# Transcriptomic analysis reveals differential regulation of synaptic components in pediatric and adult brain tumors

**DOI:** 10.64898/2026.09.30.755821

**Authors:** Leyre Merino-Galan, Sonali Arora, Gregory Glatzer, Matt Jensen, Ashmitha Rajendran, Eric C. Holland, Siobhan S. Pattwell

**Affiliations:** Ben Towne Center for Childhood Cancer and Blood Disorders Research, Seattle Children’s Research Institute, Seattle, WA, United States of America; Division of Human Biology, Fred Hutchinson Cancer Research Center, Seattle, WA, United States of America; Department of Biomedical Informatics and Education, University of Washington, Seattle, WA, United States of America; Department of Pediatrics, University of Washington School of Medicine, Seattle, WA, United States of America

## Abstract

Neuronal-cancer cell interactions are a key component of brain tumor pathophysiology, yet many fundamental questions remain unanswered – with even more remaining unasked. Here, we present a comprehensive brain tumor reference map generated from bulk RNA-seq data of ∼4,300 adult and pediatric brain tumors—including various glioma subtypes, medulloblastomas, ependymomas, and meningiomas—alongside ∼1,400 healthy brain samples. By analyzing synaptic processes and synaptic gene expression across tumor types, we reveal the complexity of neuron-tumor crosstalk and its potential impact on patient survival. Our findings identify key synaptic signaling pathways and dysregulated patterns in brain tumors, including *HNRNPH2* upregulation, a key factor in RNA processing and metabolism, in gliomas and medulloblastomas. Additionally, we highlight the role of cholinergic signaling, particularly *CHRNA9*, as a driver of glioma progression, which is also upregulated in WNT and Group 4 medulloblastomas, ependymomas, and meningiomas. Furthermore, P2 purinergic signaling emerges as a key player in glioma progression. These insights lay the groundwork for future studies exploring whether targeting these pathways could reprogram the tumor microenvironment toward an anti-tumor state, ultimately leading to novel therapeutic strategies and improved patient outcomes.

**Highlights:**

- A comprehensive **brain tumor reference map** was generated using bulk RNA-seq from ∼4,300 adult and pediatric brain tumors and ∼1,400 healthy brain samples.
- ***HNRNPH2*** is consistently upregulated across all adult and pediatric gliomas, as well as all medulloblastoma subgroups.
- **Nicotinic acetylcholine receptors (nAChRs)** are upregulated in glioblastomas and pediatric high-grade gliomas but downregulated in low-grade gliomas, underscoring their role in glioma aggressiveness.
- ***CHRNA9*** is a key driver of glioma progression and is also upregulated in medulloblastomas (WNT and Group 4), ependymomas, and meningiomas

**Graphical Abstract:** 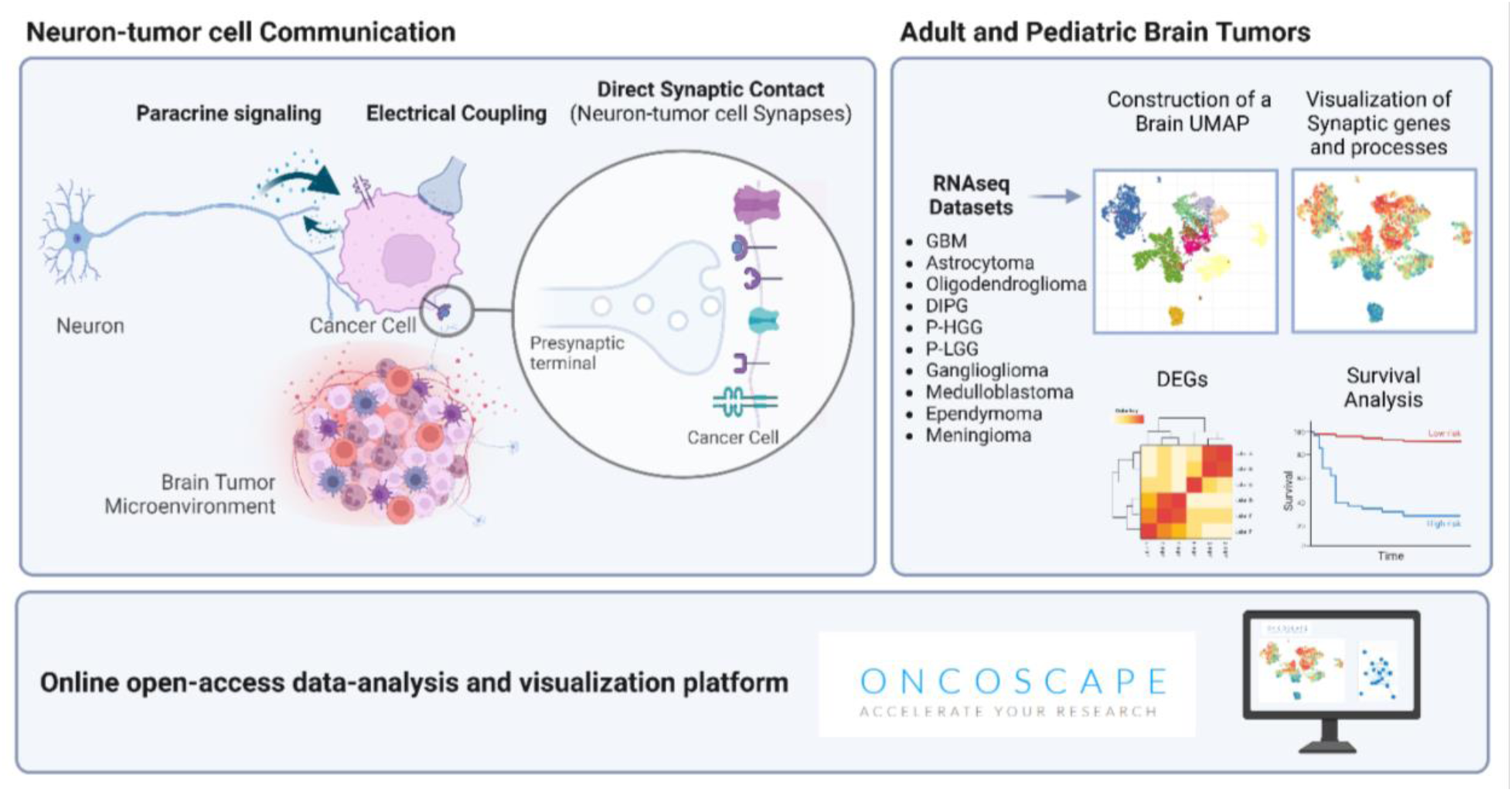

## Introduction

Brain tumors represent a clinical challenge, marked by inherent intricacies and resistance to treatment, making them among the most difficult cancers to treat. Beyond their intrinsic complexity, brain tumors are profoundly influenced by the dynamic interplay within their microenvironment, where communication between neurons and cancer cells constitutes a fundamental component of brain cancer pathophysiology^1–9^. Neurotransmitter release by surrounding neurons can profoundly impact tumor cell growth and behavior, while brain tumors, in turn, disrupt normal neurotransmitter dynamics, contributing to local neuronal dysfunction. The emerging field of cancer neuroscience research is uncovering new insights into the elaborate interplay between tumor cells and surrounding brain tissue^10,11^, yet numerous critical questions remain unanswered, underscoring the need for continued investigation in this highly synergistic and rapidly evolving field.

Neuronal activity emerges as a pivotal player in the proliferation and growth of glial malignancies. This influence is wielded through direct electrochemical synaptic communication and paracrine signaling^2–4,8^. These synaptic interactions between glioma cells and neurons, also known as neuron-glioma synapses, show typical synaptic ultrastructures, produce postsynaptic currents in glioma cells that are mediated by glutamate receptors of the AMPA subtype, and generate coordinated calcium transients that increase glioma growth and invasiveness^3,8^. Within this framework, glioma cells assume the role of a postsynaptic component, displaying morphologically diverse neuron-glioma synapses as single connections, multisynaptic contacts, or glioma cells approaching existing excitatory neuronal synapses^8^. Single-cell transcriptomic (scRNA-seq) analyses of adult and pediatric high-grade gliomas have revealed widespread and heterogeneous expression of postsynaptic receptor and scaffold genes in malignant glioma cells^3,8^, highlighting the functional relevance of synaptic biology in neoplastic brain and the surrounding normal tissue.

Beyond gliomas, various brain tumor subtypes and peripheral tumors exhibit neuron-tumor synapses and altered expression of synaptic neurotransmitters and receptors^12–15^. While research has primarily focused on the role of glutamatergic^3,8,16^ and GABAergic^6,9^ neurons in tumor pathophysiology, the influence of neuromodulatory neurons – including cholinergic, serotonergic, adrenergic, dopaminergic and purinergic neurons – remain largely unexplored. These neuronal subtypes play essential roles in normal brain function, and their dysregulation in cancer may profoundly impact tumor microenvironment (TME) interactions and neuronal activity. Although these findings underscore the importance of synaptic dynamics in tumor biology, the precise contributions of synaptic alterations across different brain tumor types remain poorly understood.

This study aims to visualize and compare the contributing role of synaptic processes and synaptic gene expression across multiple brain tumor types, and in comparison, with normal brain tissue. We combined bulk RNAseq data from 972 adult gliomas, 406 pediatric gliomas, 888 medulloblastoma, 370 ependymoma, 1,298 meningioma, 1409 samples of normal brain and 100 fetal samples and constructed a UMAP after batch correction^17–19^ to find meaningful synaptic patterns. To ensure that we ruled out batch effects, we ensured that the same bioinformatic pipeline and reference genome for each dataset. This approach enables us to compare the transcriptomic expression of synaptic proteins in brain tumors to normal brain tissue, offering a broader perspective that complements the cell-type-specific resolution provided by scRNA-seq studies. Such insights are imperative for the development of targeted therapeutic strategies aimed at disrupting the complex communication between tumor cells and their neuronal microenvironment.

## Material and Methods

### Downloading and processing bulk RNA-seq data from multiple publicly available sources

Adult glioma – Glioblastoma (GBM), astrocytoma, oligodendroglioma–, pediatric data – Diffuse intrinsic pontine glioma (DIPG), high-grade glioma (P-HGG), low-grade glioma (P-LGG) and ganglioglioma – and gene expression data for normal (control) brain samples from GTEx was obtained from Arora et al.^18^. Similarly, bulk RNA-seq data for meningioma was obtained from Thirimanne et al.^19^, while medulloblastoma and ependymoma data was obtained from Arora et al.,^17^. Batch correction was done using combatseq() function from R package “sva”^20^. After applying a variance stabilizing transformation (VST) from R package DESeq2^21^, Uniform Manifold Approximation and Project (UMAP) values were generated using the R function umap() using all protein-coding genes and visualized with R package ggplot2^22^. A 3d and 2d version of the final UMAP landscape is available via the interactive tool Oncoscape^23^.

### Differential gene expression analysis

Differential expression analysis was performed using DESeq2^21^. Significantly regulated genes in each comparison were identified based on FDR (< 0.05) and log 2fold change (> 0.3) or fold change of 25%. Batch corrected, normalized, log2 TPM counts were used to visualize the gene expression of each gene over the established reference landscape using ggplot2.

### GSVA analysis

Gene Set Variation Analysis (GSVA) ^24^ was performed on batch-corrected VST counts for all samples using Harmonizome 3.0 curated genesets^25^ and SynGO genesets^26^. The resulting GSVA scores, ranging from 1 to −1 for each sample, were visualized using ggplot2.

### Kaplan-Meier Survival Analysis

Kaplan-Meier survival curves were generated using recurrence data from tumors with known recurrence status and documented time to recurrence or last follow-up. Gene expression levels for each group were stratified into two categories: a “high” group, including patients with expression levels above the 3rd quantile, and a “low” group, comprising those with expression levels below the 1st quantile. Kaplan-Meier curves were then plotted, and p-values were calculated using the “survival” and “survminer” package in R (v3.5.7).

### Data visualization and Oncoscape integration

Matrix and clinical data were prepared for Oncoscape by converting them to cBioPortal formats (cbioportal.org). Custom settings, including colorings and precalculated views to match the paper’s figures, were stored in JSON in an Oncoscape updates.txt file. See https://github.com/FredHutch/OncoscapeV3/blob/master/docs/upload.md for details.

### Single-cell RNA Sequencing Analysis

Raw single-cell RNA sequencing data were obtained from two sources: Hovestadt et al. 2019^27^ (GSE119926), comprising 23 diagnostic medulloblastoma samples, 2 recurrence samples, and 11 patient-derived xenografts; and Abdelfattah et al. 2022^28^ (GSE182109), comprising 44 glioma samples including 3 low-grade gliomas, 17 newly diagnosed glioblastomas, and 24 recurrent glioblastomas. Individual Seurat objects were created for each sample using standard Seurat pipelines^29^. For the medulloblastoma dataset, cell cycle scoring was performed using CellCycleScoring function with canonical S-phase and G2M-phase gene sets^29,30^. For both datasets, mitochondrial gene percentages were calculated using the pattern “^MT-”, and data were normalized using SCTransform^31^ with all genes retained. Principal component analysis was performed for all samples, followed by UMAP and t-SNE dimensionality reduction. Sample integration was performed using distinct approaches for each dataset: medulloblastoma samples were integrated using the mutual nearest neighbor approach via RunFastMNN^32^ on the first 30 MNN-corrected dimensions, while glioma samples were integrated using Seurat’s anchor-based integration workflow with a k.weight parameter of 36 and subsequent PCA recomputed on 30 principal components. Graph-based clustering was performed using the FindNeighbors and FindClusters functions3 with Louvain algorithm implementation at resolutions of 0.8 (medulloblastoma) and 0.5 (glioma).

## Results

### The Brain-UMAP reveals differential dysregulation of synaptic pathways across brain tumor types

To unravel the biological intricacies of synaptic contributions to brain tumor physiology, we analyzed RNA-seq data from 28 datasets to generate a comprehensive Brain-UMAP The collection of brain tumors analyzed included neoplastic brain, adult glioblastoma (comprising oligodendroglioma, astrocytoma and GBM) and diverse pediatric brain tumors including medulloblastoma, meningioma, pediatric high-grade glioma, DIPG (**Figure 1A, B, Suppl. Figure 1A, Supp Table 1**).

**Figure 1:**
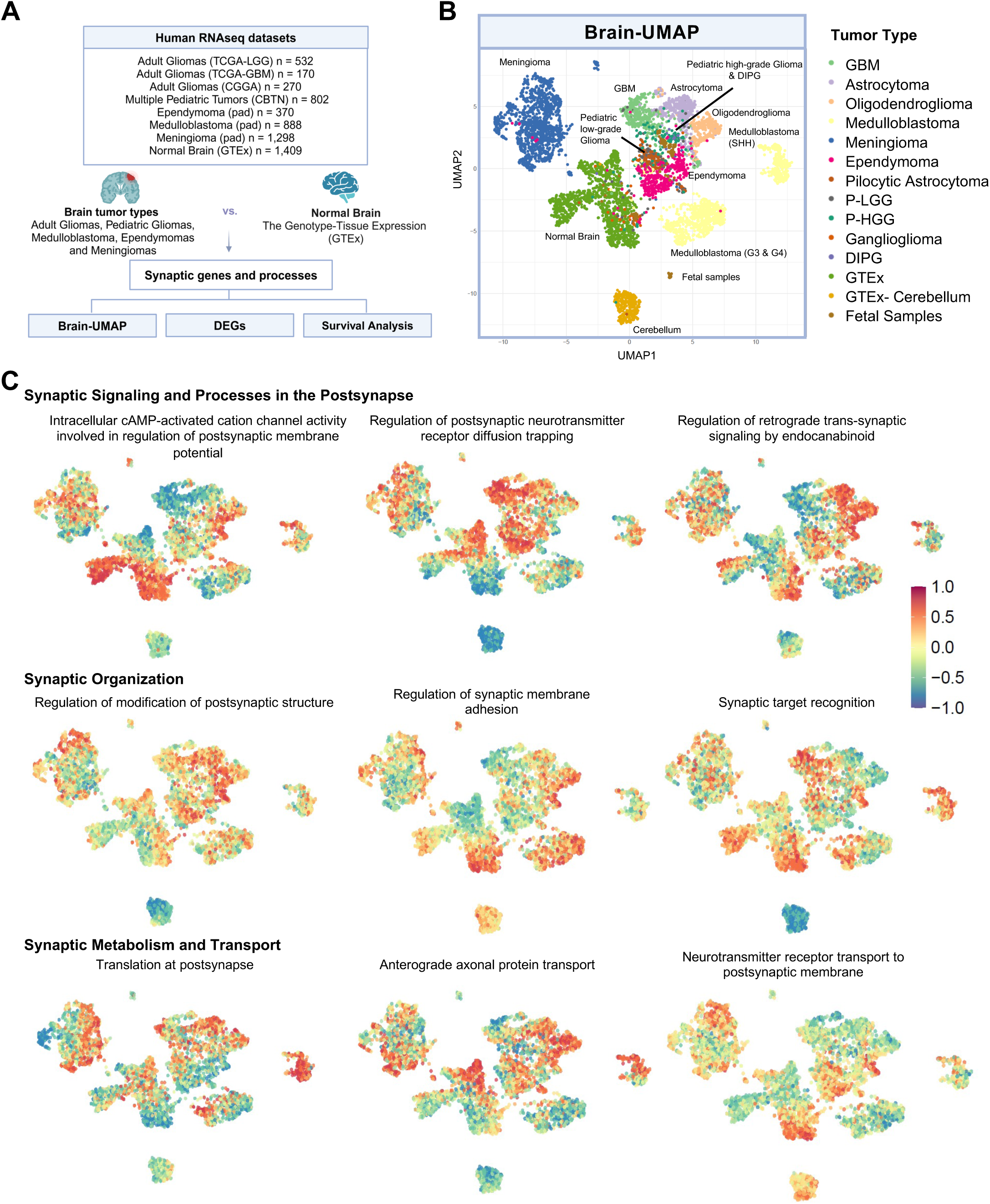
Synaptic Process Involvement Across Brain Tumor Types. **(A)** Overview of the data analysis pipeline, including brain tumor types, RNA-seq datasets used, and the analytical steps: selection of synaptic genes and related processes from SynGO dataset, construction of the Brain-UMAP, differential gene expression analysis (DEGs), and survival analysis. **(b)** UMAP representation of the full dataset, including adult gliomas from TCGA (The Cancer Genome Atlas Low Grade Glioma) and CGGA (Chinese Glioma Genome Atlas); pediatric tumors from CBTN (Children’s Brain Tumor Network); medulloblastomas, ependymomas, and meningiomas from publicly available datasets (pad); and normal brain and cerebellum samples from GTEx (Genotype-Tissue Expression project). **(C)** Brain-UMAPs highlighting SynGO-derived gene ontology (GO) terms related to synaptic signaling and postsynaptic processes, synaptic organization, and synaptic metabolism and transport. CBTN, Children’s Brain Tumor Network; CGGA, Chinese Glioma Genome Atlas; DEG, differential gene expression analysis; DIPG, diffuse intrinsic pontine glioma; GBM, glioblastoma; GTEx, Genotype-Tissue Expression project; *Pad*, publicly available dataset; P-LGG, pediatric low-grade glioma; P-HGG, pediatric high- grade glioma; TCGA, The Cancer Genome Atlas Low Grade Glioma.

This detailed gene expression landscape sheds light on the dysregulation of synaptic signaling, organization, metabolism, and transport pathways across various cancer types in comparison to normal brain tissue. Using GSVA with SynGO pathway annotations, we identified distinct patterns of synaptic pathway enrichment across tumor subtypes (**Figure 1C**). For instance, aggressive GBM displayed upregulation in pathways such as “Postsynaptic neurotransmitter diffusion trapping” and “Synaptic target regulation,” patterns that were similarly observed in P-HGG and DIPG. Conversely, astrocytomas, oligodendrogliomas, and P-LGG showed enrichment in pathways associated with “Regulation of retrograde trans-synaptic signaling by endocannabinoids” and “Anterograde axonal protein transport.“

Beyond gliomas, other tumor types exhibited subtype-specific synaptic alterations. Medulloblastomas demonstrated enhanced activity in pathways like “Synaptic membrane adhesion,” “Synaptic target recognition,” and “Translation at the postsynapse,” while ependymomas showed upregulation in “Postsynaptic receptor diffusion trapping”. Meningiomas presented a broader enrichment of synaptic processes, with notable increases across multiple pathways (**Figure 1C**). These findings underscore the tumor-specific nature of synaptic alterations, emphasizing the need for further investigation into their functional implications. All SynGO pathways and genes mapped onto the Brain-UMAP are accessible via the Oncoscape webpage (https://github.com/sonali-bioc/ComprehensiveBrainUMAP).

### The brain tumor landscape reveals distinct neuronal signatures across various brain cancer types

Given that specific neuronal subtypes innervate different brain regions and tumors, exerting diverse functions that influence both cancer cells and the TME, we analyzed the chemical synaptic signaling profiles associated with these neuronal signatures (**Figure 2A**). To date, research efforts have mainly focused on the influence of glutamatergic ^3,8,16^ and GABAergic ^6,9^ neurons on tumor pathophysiology, while the effects of neuromodulatory neuron types - cholinergic, serotonergic, adrenergic, dopaminergic and purinergic neurons - remain largely unexplored. Our study focuses on synaptic receptor expression because neurotransmitter gene transcription primarily occurs in the soma, while synaptic terminals, where neurotransmitter signaling takes place, may be located far from the site of transcription. In the context of tumors, where bulk tissue analysis is performed, this spatial separation limits the ability to capture neurotransmitter dynamics. Instead, receptor expression serves as a more reliable marker of synaptic signaling alterations, as local regulatory mechanisms fine-tune receptor expression and function ^33^. Our findings reveal a differential expression of specific synaptic receptor subtypes (**Figure 2B, Supp Table 2**) and receptor groups (**Suppl. Figure 2A**) across various brain tumor types. For instance, we identified elevated expression of glutamatergic receptors in both adult and pediatric low-grade gliomas, whereas GBM and DIPG exhibited high expression of specific cholinergic receptors. Interestingly, gangliogliomas demonstrated elevated expression across multiple synaptic receptor types. In medulloblastoma subtypes, distinct receptor profiles emerged: Group 3 tumors showed high expression of GABRR receptors, Group 4 displayed elevated muscarinic and serotonin receptor expression, and SHH tumors exhibited high expression of nicotinic receptors.

**Figure 2:**
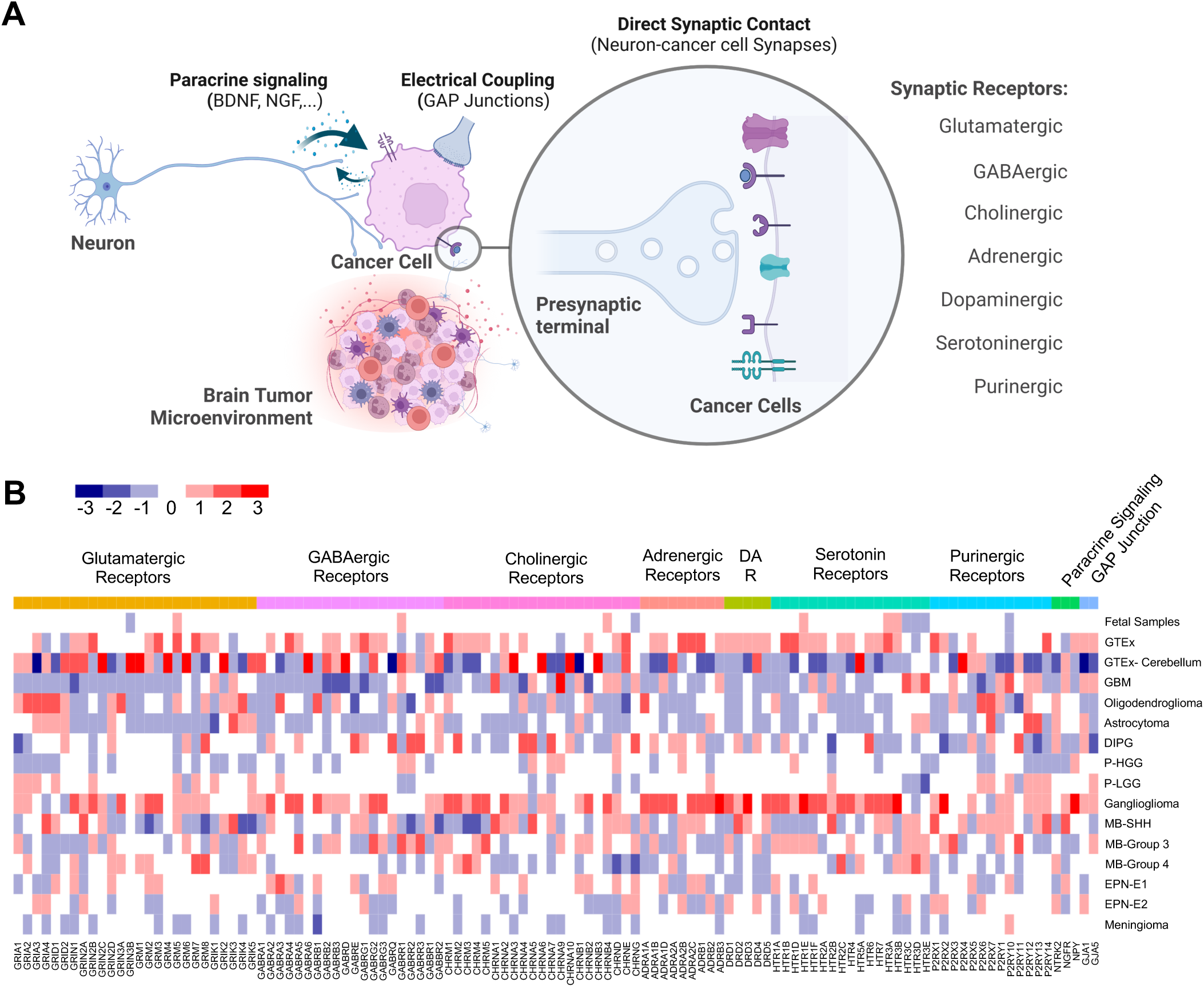
Neuronal Regulation of Brain Tumors. **(A)** Schematic representation of the proposed mechanisms of communication between neurons and brain tumor cells, including paracrine signaling, electrical coupling via gap junctions, and direct synaptic contact in which a presynaptic neuron forms a synapse with a cancer cell expressing postsynaptic receptors. **(B)** Transcriptomic analysis reveals differential enrichment of synaptic receptor genes across various brain tumor types. The analysis includes glutamatergic, GABAergic, cholinergic, adrenergic, dopaminergic (DA), serotonergic, and purinergic receptors, as well as paracrine signaling molecules and receptors (*NTRK2*, *NGFR*, *NPY*) and gap junction proteins (*GJA1*, *GJA5*). Synaptic gene expression was compared across tumor types, and enrichment scores indicate upregulation (positive values) or downregulation (negative values). DA R, dopaminergic receptors; DIPG, diffuse intrinsic pontine glioma; EPN, ependymoma; GBM, glioblastoma; GTEx, Genotype-Tissue Expression project; LGG, low-grade glioma; MB, medulloblastoma; MB-SHH, sonic hedgehog group.

While direct chemical synaptic signaling remains the best-characterized form of neuronal communication, neurons and cancer cells also interact through alternative mechanisms, including paracrine signaling via neurotrophic factors (brain derived neurotrophic factor (BDNF), nerve growth factor (NGF)) and neuropeptides (neuropeptide Y (NPY)), as well as electrical coupling ^2–4^. Our analysis revealed distinct patterns of expression for key receptors involved in these pathways (**Figure 2B** and **Suppl. Figure 2B**). *NGFR* was highly significantly expressed in SHH medulloblastoma(log2foldchange = 1.59, adj-pvalue = 1.24975851071571E-28), ganglioglioma (log2foldchange = 0.8099, adj-pval = 0.01) and GBM (log2foldchange= 1.058184, adj-pval = 1.915202e-18), while *NTRK2* expression was elevated in astrocytoma, oligodendroglioma, pilocytic astrocytoma, and Group 4 medulloblastoma. Notably, *NPY* expression was particularly enriched in ganglioglioma and pediatric high-grade gliomas. Regarding electrical coupling, *GJA1* (Connexin 43) was highly expressed in GBM, astrocytoma, ependymoma (EP-E2) and ganglioglioma (**Figure 2B** and **Supp. Figure 2C**). These findings underscore the diverse non-direct synaptic interactions shaping tumor-neuron communication across brain malignancies.

### Adult gliomas exhibit shared and subtype-specific synaptic gene alterations of GABAergic, glutamatergic, cholinergic, and purinergic pathways with distinct prognostic implications

Glioma cells are known to form synaptic-like structures with neurons assuming roles analogous to postsynaptic components^3,8^. These interactions enhance glioma cell proliferation and invasion^3,8^, highlighting the importance of synaptic machinery in tumor progression. To investigate this further, we conducted a differential gene expression analysis across three distinct adult glioma types, comparing their expression profiles to normal brain tissue (**Figure 3A, B, Supp Table 3**). Our analysis assessed both upregulated and downregulated DEGs that overlap with SynGO-annotated genes. In GBM, we identified 19 upregulated and 56 downregulated synaptic genes, while oligodendroglioma displayed 22 upregulated and 46 downregulated synaptic genes, and astrocytoma showed 4 upregulated and 31 downregulated synaptic genes (**Figure 3B**). These DEGs encompass various postsynaptic receptor types, ion channels, scaffold proteins, cytoskeletal elements, and synaptic regulators (**Figure 3C, D**). Interestingly, we identified four synaptic genes that were consistently upregulated across all three glioma subtypes*: RPL36A, HNRNPH2, ATP6V0C, and VAMP2* **(Figure 3C, F)**. Notably, VAMP2 is a key component of the SNARE complex, facilitating synaptic vesicle docking and fusion at the presynaptic membrane. ATP6V0C is part of the vacuolar ATPase, which regulates intracellular and extracellular pH by pumping protons into vesicles and the extracellular environment, and HNRNPH2 is an RNA-binding proteins involved in mRNA processing, splicing, transport, and stability. In contrast, six synaptic genes were consistently downregulated across the glioma subtypes: *FGA, CPLX3, SLC18A2, GRP, SLC6A3*, and *DRD3*, primarily involved in monoamine and dopamine transport (**Figure 3D, Supplementary Figure 3C**). The SynGO gene ontology analysis further revealed that the differentially expressed genes were strongly associated with specific synaptic components and processes (**Suppl. Figure 3A, B**).

**Figure 3:**
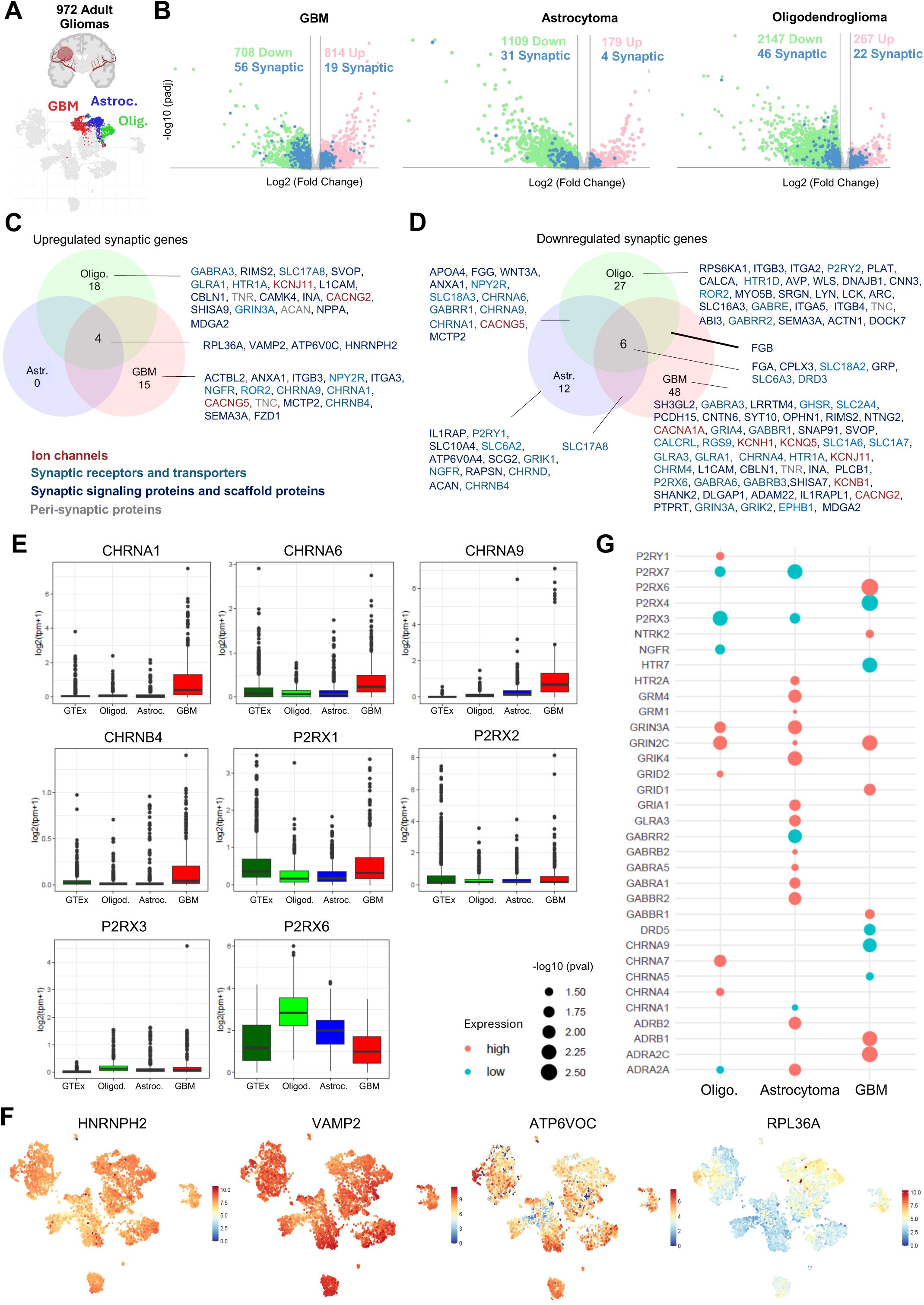
Synaptic Gene Dysregulation Across Adult Glioma Subtypes. **(A)** Brain-UMAP representation highlighting glioblastoma (GBM, red), astrocytoma (blue), and oligodendroglioma (green). **(B)** Volcano plots showing differentially expressed genes (DEGs) for GBM, astrocytoma, and oligodendroglioma compared to normal brain tissue (GTEx). Upregulated genes are shown in pink, downregulated genes in green, and differentially expressed synaptic genes are indicated in blue. **(C, D)** Venn diagrams comparing shared upregulated (**C**) and downregulated (**D**) genes among the three adult glioma types. Genes within each cluster are color-coded by synaptic function: ion channels (dark red), synaptic receptors and transporters (light blue), synaptic signaling and scaffold proteins (dark blue), and peri-synaptic proteins (gray). **(E)** Box plots showing log₂ gene expression of selected cholinergic (*CHRNA1*, *CHRNA6*, *CHRNA9*, *CHRNB4*) and purinergic (*P2RX1*, *P2RX2*, *P2RX4*, *P2RX6*) receptors in normal brain (GTEx), oligodendroglioma, astrocytoma, and GBM. **(F)** Expression patterns of shared upregulated synaptic genes (*HNRNPH2*, *VAMP2*, *ATP6V0C*, *RPL36A*) for the three adult glioma types visualized across the Brain-UMAP. **(G)** Dot plot showing the association between synaptic receptor gene expression and survival in oligodendroglioma, astrocytoma, and GBM. Only synaptic genes with statistically significant p-values are displayed (*p* < 0.05). Genes whose higher expression is associated with poorer survival are shown in orange, while those linked to better prognosis are shown in turquoise. Dot size represents the statistical significance of the association, expressed as –log₁₀(p-value). GBM, glioblastoma; GTEx, Genotype-Tissue Expression project

We next investigated the molecular pathways of the synaptic DEGs. Our findings revealed distinct patterns of synaptic gene expression, highlighting glioma-specific adaptations to the neural microenvironment. In GBM, nicotinic acetylcholine receptor subunits (*CHRNA1, CHRNB4, CHRNA9*) and a neurotrophic growth factor receptor (*NGFR*) were highly expressed (**Figure 3C**). In contrast, key synaptic components such as GABA receptor subunits (*GABRA6, GABRB3, GABRA3, GABBR1*), ionotropic glutamate receptors (*GRIA4, GRIN3A, GRIK2*), monoamine receptors (*HTR1A, CHRM4*) and peptide secretion regulators (*PLCB1, GHSR, GRP*) showed reduced expression (**Figure 3D**). The downregulation of GABA receptors along with suppressed glutamate and monoamine receptor signaling, suggests widespread dysregulation of both inhibitory and excitatory neurotransmission. These alterations may disrupt synaptic homeostasis, fostering a tumor-promoting microenvironment and contributing to GBM’s aggressive phenotype.

Oligodendrogliomas exhibited increased expression of specific glutamatergic (*GRIN3A*), GABAergic (*GABRA3*) and serotoninergic (*HTR1A*) receptors, alongside modulators of synaptic transmission (*SHISA9, CACNG2*) (**Figure 3C**). Conversely, nicotinic acetylcholine receptor subunits (*CHRNA1, CHRNA6, CHRNA9*), GABA receptor subunits (*GABRR2, GABRR1, GABRE*) and interestingly, focal adhesion molecules (*ITGB4, TNC, ITGA5, ITGA2, ACTN1, ITGB3*) were consistently downregulated, suggesting decreased synaptic connectivity and cellular adhesion (**Figure 3D**). In astrocytoma, synaptic receptor expression profiles largely resembled those of normal brain tissue with a few exemptions, such as the downregulation of nicotinic acetylcholine receptors (*CHRNA1, CHRNB4, CHRNA6, CHRNA9*) and specific glutamate (*GRIK10*) and GABA (*GABRR*) receptor subunits (**Figure 3C, D**). This may indicate a less aggressive adaptation to the microenvironment. Notably, purinergic receptor subtypes, which are versatile regulators of synaptic function, were downregulated in all glioma types (*P2RX6* in GBM, *P2RY1* in astrocytoma and *P2RY2* in oligodendroglioma) (**Figure 3E**). Importantly, the differential expression of nicotinic acetylcholine receptors (**nAChRs**)—upregulated in GBM but downregulated in low-grade gliomas (**Figure 3E**)—underscores their potential role in glioma aggressiveness.

Survival analysis revealed distinct associations between specific synaptic genes and prognosis across glioma subtypes. In GBM, high expression levels of *P2RX6, NTRK2, GRIN2C, GRID1, GABBR1, ADRB1 and ADRA2C* receptors were linked to poor prognosis (**Figure 3G**). In contrast, *CHRNA9 and CHRNA5* low expression was associated with poor prognosis. Given its prominent role in glioma aggressiveness, nAChR subunits represent promising therapeutic targets for disrupting these tumor-promoting pathways and restoring a more inhibitory or homeostatic environment. In contrast, poor prognosis in astrocytoma was associated with high expression *of HTR2A, GRM4, GRM1, GRIN3A, GRIN2C, GRIK4, GRIA1, GLRA3, GABRB2, GABRA1, GABRA5, GABBR2, ADRB2 and ADRA2A,* whereas oligodendroglioma showed poor prognosis linked to high expression of *P2RY1, GRIN3A, GRIN2C, GRID2, CHRNA7, CHRNA4* (**Figure 3G**).

### Pediatric glioma types show subtype-specific synaptic gene dysregulation in GABAergic, glutamatergic, and cholinergic pathways

Neuronal activity also drives the progression of both pediatric high-grade ^2^ and low-grade gliomas^5^ facilitated through paracrine signaling and direct glutamatergic and GABAergic neuron-glioma synapses ^2,3,5,6,9^. Our differential expression analysis revealed distinct patterns of synaptic gene regulation across pediatric glioma subtypes. In DIPG, 20 genes were upregulated and 96 were downregulated. P-HGGs exhibited 10 upregulated and 50 downregulated synaptic genes, while P-LGGs displayed 8 upregulated and 73 downregulated synaptic genes. Gangliogliomas showed a markedly different profile, with 153 upregulated and 34 downregulated synaptic genes (**Figure 4B, Supp Table 4**). Among these, *HNRNPH2* was the only synaptic gene consistently upregulated across all four pediatric glioma types (**Figure 4C**), mirroring its upregulation in adult gliomas (**Figure 3F**). Although not exclusively linked to synaptic function, HNRNPH2 plays a critical role in RNA metabolism, with recent evidence highlighting its involvement in neuronal and synaptic regulation^34^. Mutations in HNRNPH2 are associated with X-linked syndromic intellectual disability^35^, often presenting with synaptic dysfunction, further underscoring its importance in synaptic biology. In contrast, 13 synaptic genes were consistently downregulated across all pediatric glioma types, suggesting impaired neurotransmitter transport (*SLC22A2, SLC6A3, SLC1A7, SLC6A50),* synaptic receptor function (*GRIN3B, GRM4, GABRA6, CHRNB3, GLRA1*), and synapse-associated signaling (*CBLN1, WNT3A, CLEC3A, FGB*) (**Figure 4D**). This widespread suppression of glutamatergic, GABAergic, and cholinergic pathways may contribute to tumor-associated neurodevelopmental alterations and the overall impact of gliomas on pediatric brain function.

**Figure 4:**
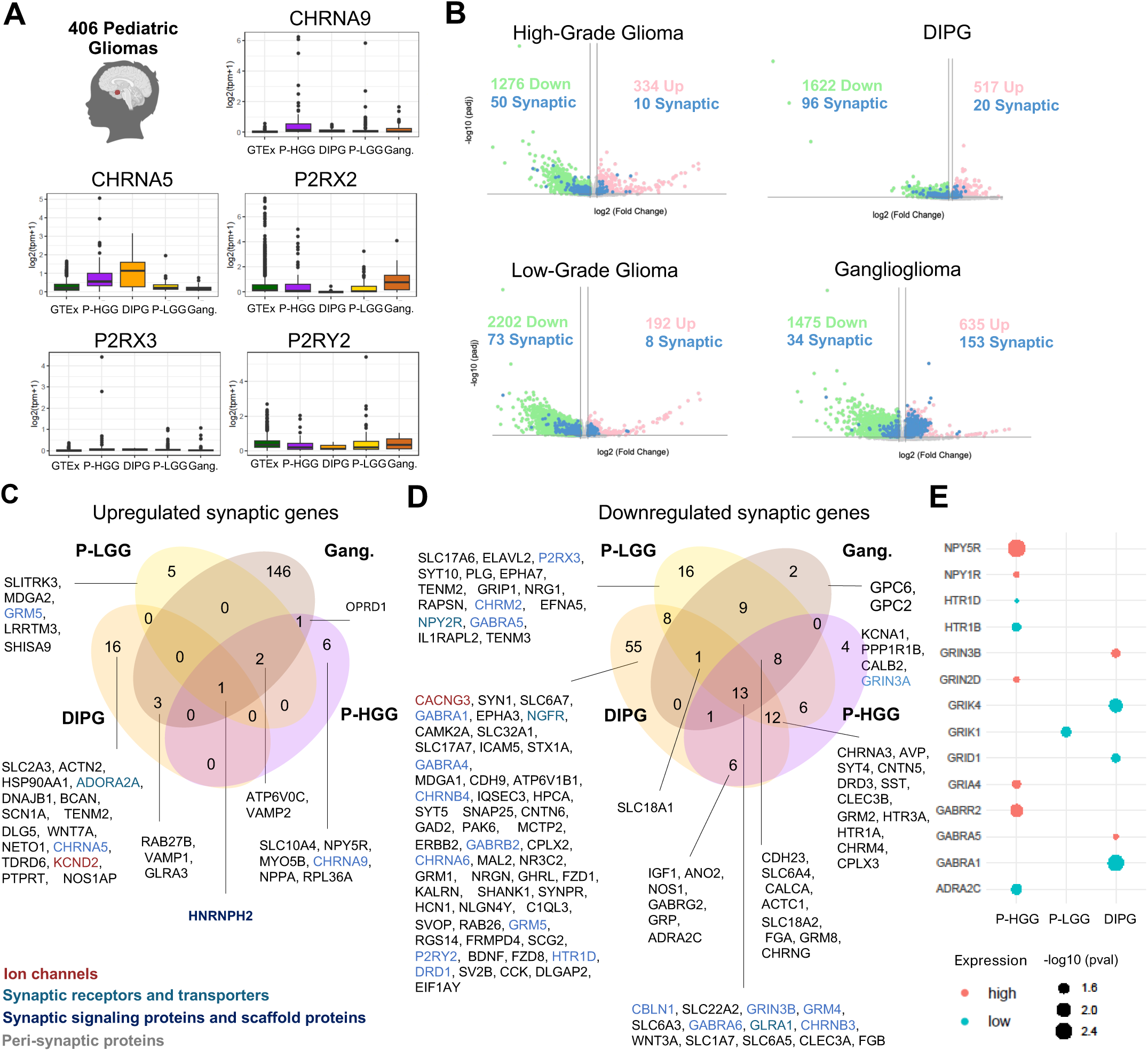
Differential Expression and Enrichment of Synaptic Genes in Pediatric Gliomas. **(A)** Box plots showing log₂ gene expression of selected cholinergic (*CHRNA9*, *CHRNA5*) and purinergic (*P2RX2*, *P2RX3*, *P2RY2*) receptors in normal brain (GTEx), pediatric high-grade gliomas (P-HGG), DIPG, pediatric low-grade gliomas (P-LGG) and gangliogliomas. **(B)** Volcano plots showing differentially expressed genes (DEGs) for pediatric high-grade gliomas, DIPG, pediatric low-grade gliomas and gangliogliomas compared to normal brain tissue (GTEx). Upregulated genes are shown in pink, downregulated genes in green, and differentially expressed synaptic genes are indicated in blue. **(C, D)** Venn diagrams comparing shared upregulated (**C**) and downregulated (**D**) genes among the four pediatric glioma types. Genes within each cluster are color-coded by synaptic function: ion channels (dark red), synaptic receptors and transporters (light blue), synaptic signaling and scaffold proteins (dark blue), and peri-synaptic proteins (gray). **(E)** Dot plot showing the association between synaptic receptor gene expression and survival in P-HGG, DIPG, P-LGG and gangliogliomas. Only synaptic genes with statistically significant p-values are displayed (*p* < 0.05). Genes whose higher expression is associated with poorer survival are shown in orange, while those linked to better prognosis are shown in turquoise. Dot size represents the statistical significance of the association, expressed as –log₁₀(p-value).

Pediatric high-grade gliomas shared molecular similarities with their adult counterparts. Nicotinic cholinergic receptor subunits were upregulated, with *CHRNA9* elevated in P-HGGs and *CHRNA5* in DIPG (**Figure 4A**). In contrast, purinergic receptors were consistently downregulated, including *P2RY2* in DIPG, *P2RX3* in P-LGGs, and *P2RX2* across both pediatric glioma types (**Figure 4A**). DIPG exhibited widespread downregulation of synaptic genes, affecting multiple neurotransmitter systems. This included nicotinic acetylcholine receptor subunits (*CHRNA1, CHRNA3, CHRNA6, CHRNB3, CHRNB4*), GABA receptor subunits (*GABRA1, GABRA4, GABRA6, GABRB2, GABRG2*), glutamate receptors (*GRIN3B, GRM2, GRM4*), and serotonin receptors (*HTR1A, HTR1D, HTR3A*) (**Figure 4D**). Additionally, several key regulators of trans-synaptic signaling were downregulated (*BDNF, CAMK2A, ERBB2, NOS1, CACNG3, STX1A, WNT3A, FZD8, ATP6V1B1, SHANK1, DLGAP2* and *NLGN4Y*), while other P-HGGs also exhibited downregulation of the glutamatergic receptor *GRIN3A* and dopaminergic receptors *DRD2* and *DRD3* (**Figure 4D**). These findings suggest that high-grade gliomas, particularly DIPG, undergo extensive synaptic impairments, which may contribute to their aggressive behavior and resistance to treatment.

In contrast, P-LGGs exhibited a more limited impact on synaptic gene networks, which may correlate with their less invasive nature. This was characterized by the upregulation of *GRM5*, a metabotropic glutamate receptor, while *CHRM2*, *GABRA5*, and *NPY2R* were downregulated (**Figure 4C, D**). Notably, *GABRR1*, *GABRR2*, and *CHRNA5* were downregulated in both P-LGGs and gangliogliomas, suggesting some shared synaptic alterations in GABAergic and cholinergic signaling in these tumor types.

Gangliogliomas, predominantly benign and slow-growing tumors, are defined by their dual neuronal and glial composition. Unsurprisingly, these tumors showed a significant upregulation of synaptic genes compared to normal brain tissue (**Figure 4C**). Specifically, gangliogliomas demonstrated enhanced expression across several key pathways including GABA receptor related genes (*GAD1, GAD2*, *GABRA4, GABRB2, GABRB1, GABBR1, GABRA5, GABRA1, GABRA2, GABRG3*), NMDA receptor activation and long-term potentiation (*APBA1, NRGN, LRRC7, GRIN2A, GRIN3A, GRIN1, DLG2, CAMK2B, PRKAR1B, GRIN2B, CAMK2A*) and neurotransmitter release cycle (*SLC32A1, SLC17A7, RAB3A, STX1A, SLC6A11, SNAP25, SYT1, APBA1, SLC1A2, SYN1, CPLX1, GAD1, GAD2*). The extensive involvement of pathways such as GABAergic and glutamatergic signaling and synaptic plasticity underscores the role of gangliogliomas in maintaining a robust neuronal network within the TME.

Despite the large number of DEGs identified in ganglioglioma, none were associated with changes in survival. Only *GRIK1* low expression correlated with poorer prognosis in low-grade gliomas. In contrast, a distinct pattern emerged in pediatric high-grade gliomas, where high expression of *GRIA4, GABRR2, NPY1R and NPY5R* was linked to unfavorable outcomes, while in DIPG high expression of *GRIN3B*, and *GABRA5* was associated with the worst prognosis **(Figure 4E).**

### Subgroup-Specific Synaptic Gene Deregulation Highlights Neurobiological Complexity in Medulloblastoma

Synaptic and neuronal gene upregulation has been identified in specific medulloblastoma subgroups, particularly in SHH^36–38^. These observations suggest that medulloblastoma cells might adopt neuronal-like properties or exploit synaptic machinery, similar to what has been observed in gliomas^3,8^. To investigate this further, we performed a DEG analysis across the four medulloblastoma subgroups, comparing them to normal cerebellar tissue, to determine the extent and specificity of synaptic pathway alterations (**Figure 5A, Supp Table 5**). While the use of adult cerebellum as a control may not be ideal due to its mature nature, it remains the anatomically most appropriate comparison for our study.

**Figure 5:**
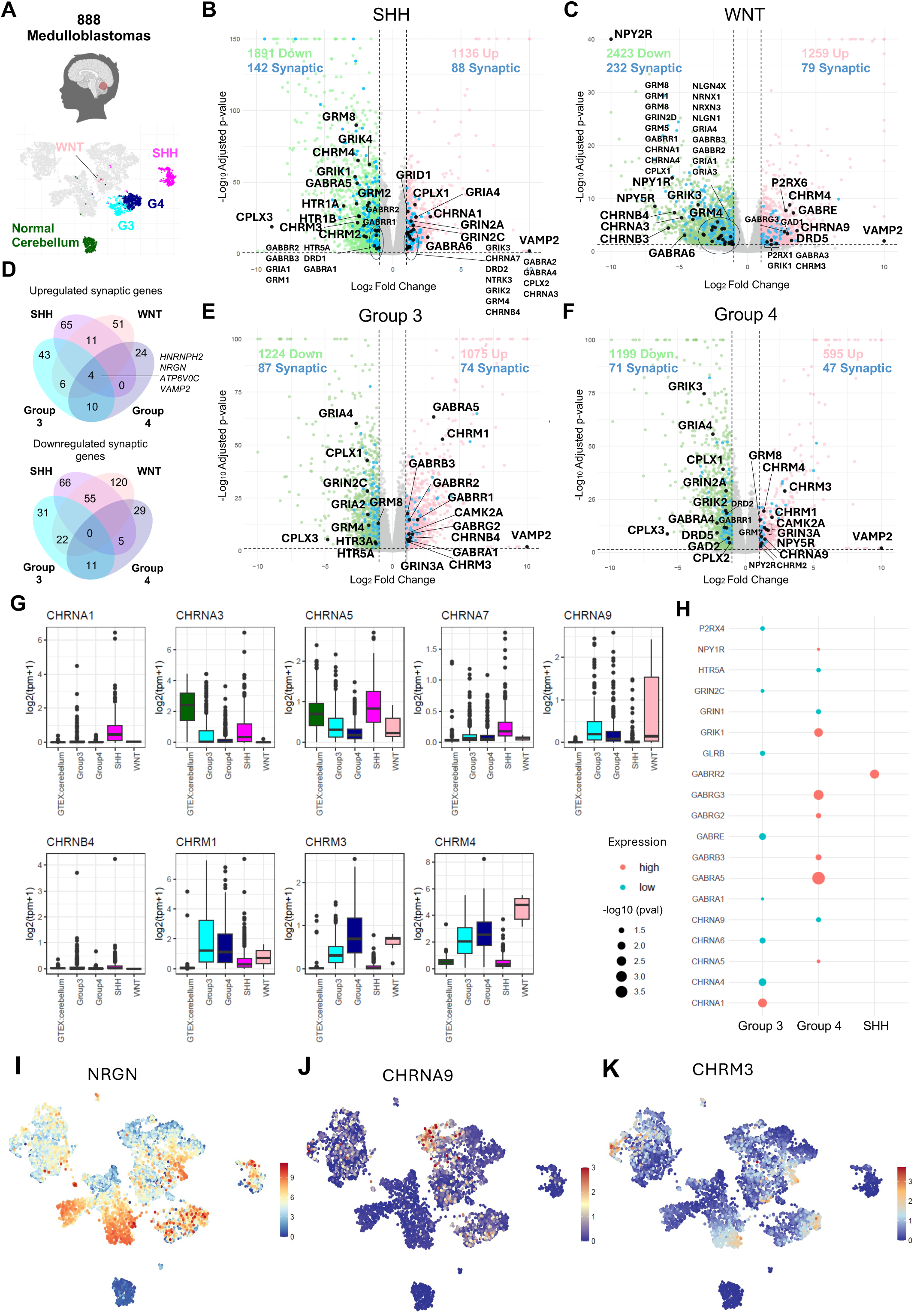
Subtype-Specific Synaptic Gene Dysregulation in Medulloblastoma Compared to Normal Cerebellum. **(A)** Brain-UMAP representation highlighting medulloblastoma subgroups: SHH (purple), Group 3 (light blue), Group 4 (dark blue) and WNT (light pink). **(B, C, E, F)** Volcano plots showing differentially expressed genes (DEGs) for medulloblastoma subgroups: (**B**) SHH, (**C**) WNT, (**D**) Group 3 and (**E**) Group 4 compared to cerebellum tissue (GTEx). Upregulated genes are shown in pink, downregulated genes in green, and differentially expressed synaptic genes are indicated in blue or black (highlighted). **(D)** Venn diagrams comparing shared upregulated and downregulated genes among the four medulloblastoma subgroups. **(G)** Box plots showing log₂ gene expression of selected cholinergic receptors (*CHRNA1*, *CHRNA3*, *CHRNA5, CHRNA7, CHRNA9*, *CHRNB4, CHRM1, CHRM3, CHRM4*) in normal cerebellum (GTEx) and medulloblastoma subgroups. **(H)** Dot plot showing the association between synaptic receptor gene expression and survival in medulloblastoma subgroups: Group 3, Group 4 and SHH. Only synaptic genes with statistically significant p-values are displayed (*p* < 0.05). Genes whose higher expression is associated with poorer survival are shown in orange, while those linked to better prognosis are shown in turquoise. Dot size represents the statistical significance of the association, expressed as –log₁₀(p-value). **(I, J, K)** Expression patterns of (**I**) *NRGN*, (**J**) *CHRNA9* and (**K**) *CHRM3* visualized across the Brain-UMAP.

Our analysis revealed a significant deregulation of synaptic proteins across all four medulloblastoma subgroups (**Figure 5B-F**). These proteins, critical to pre- and post-synaptic membranes and chemical synaptic transmission (**Suppl. Figure 5A, B**), showed distinct expression patterns. Among the subgroups, the SHH group exhibited the highest number of upregulated synaptic proteins (88), followed by 79 in the WNT group, 74 in Group 3, and 47 in Group 4 (**Figure 5B-F**). Notably, all subgroups shared four upregulated synaptic genes (*HNRNPH2, NRGN, ATP6V0C, and VAMP2*) (**Figure 5D**), which were also upregulated in adult gliomas. *NRGN* (neurogranin) is a neuronal calmodulin-binding protein crucial for synaptic plasticity. *NRGN* upregulation could reflect a shift towards neuronal-like properties, affecting tumor plasticity and neuron-tumor interactions (**Figure 5I**). In contrast, the downregulated synaptic genes varied more significantly across subgroups. The WNT group demonstrated the highest number of downregulated synaptic genes (232), followed by 142 in SHH, 87 in Group 3, and 71 in Group 4 (**Figure 5B-F**). Interestingly, no synaptic genes were commonly downregulated across all subgroups (**Figure 5D**).

Focusing on the SHH subgroup, we observed an upregulation of genes involved in various neurotransmission and signaling pathways that include metabotropic (*GRM4*) and ionotropic glutamatergic signaling (*GRIA4, GRIN2A, GRIN2C, GRIK2, GRIK3, GRID1*), nicotinic cholinergic signaling (*CHRNA1, CHRNA3, CHRNA5, CHRNA7, CHRNB4*), neurotransmitter release cycle (*VAMP2, CPLX1, CPLX2, SYT1, ATP6V0C, SLC18A1, SLC18A2, SLC18A3, SLC18A8*), GABAergic signaling (*GAD2, GABRA2, GABRA4, GABRA6*) and neurotrophic signaling (*NGFR, NTRK3, MAPK10*) and *DRD2* (**Figure 5B**). Conversely, significant downregulation in the SHH subgroup was noted in genes associated with glutamatergic signaling (*GRIA1, GRIN3A, GRIK1, GRIK4, GRM1, GRM2, GRM7, GRM8, PRKCG, CACNA1C, ADCY1, SHANK1, DLGAP1*), GABAergic and dopaminergic synapses (*GABBR2, GABRB3, GABRA5, GABRR2, GABRA1, GABRR1, DRD1*), serotoninergic synapses (*HTR1A, HTR5A*, *HTR1B),* cholinergic signaling *(CHRM3, CHRM2, CHRM4*, *CHRNG)* and synaptic vesicle cycle *(SLC32A1, SLC17A7, SLC6A7, SLC6A4, SLC6A3, STX3, CPLX3, SLC6A5)* (**Figure 5B**). Despite its overall downregulation in SHH medulloblastomas, higher expression of *GABRR2* correlated with worst prognosis, suggesting that *GABRR2* may serve as a prognostic marker (**Figure 5H**).

In the WNT subgroup, upregulated genes were associated with several pathways, including GABAergic signaling (*GAD1, GABRA3, GABRE, GABRG3*), cholinergic signaling (*CHRM3, CHRM4, CHRNA9, PLCB1, PLCB2, PLCB3*), purinergic signaling (*P2RX1, P2RX6*), synaptic vesicle cycle (*STX1A, ATP6V0C, STX3, VAMP2, CACNA1A)* and *DRD5 and GRIK1* (**Figure 5C**). The WNT subgroup also exhibited downregulation in pathways such as glutamatergic signaling (*GRIN2D, GRIA4, GRM1, GRIA2, GRM5, HOMER1, GRM8, GRM3, GRIK3, GRIA1, GRM4, GRIA3*), GABAergic synapse (*GABBR2, GABRA6, GABRB1, GABRB3, GAD2, GABRA5, GABRA1, GABRR1*), nicotinic acetylcholine receptors (*CHRNA1, CHRNA3, CHRNA4, CHRNB3, CHRNB4*), synaptic vesicle cycle (*STX1B, SLC32A1, SLC1A6, RAB3A, SLC6A7, SLC6A4, SLC1A1, SLC6A3, CPLX1, ATP6V0A4, DNM3, CACNA1B, RIMS1*), neuropeptide Y signaling (*NPY1R, NPY2R, NPY5R*), glycine receptors (*GLRA2, GLRA3, GLRB*) and neurexins and neuroligins (*NLGN4X, NRXN1, NLGN1, NRXN3*). While NLGN3 promotes tumor proliferation via activity-dependent mechanisms in gliomas^2,4^, the downregulation of NLGNs and NRXNs, key molecules involved in adhesion, synapse formation, and maintenance, along with a broad range of synaptic receptors in WNT medulloblastoma suggests that these tumors do not rely on synaptic signaling for progression. Instead, this downregulation may indicate weakened cell-cell and synaptic interactions, potentially impacting tumor migration, invasion, or adhesion within the cerebellum. Although the small sample size precluded survival analysis, these findings underscore the unique and dualistic synaptic landscape of the WNT subgroup, shedding light on its distinct synaptic characteristics.

Group 3, the medulloblastoma group with the poorest survival rate among the medulloblastoma subtypes, showed upregulation in GABAergic receptor signaling (*GABRB3, GABRA5, GABRR2, GABRA1, GABRR1, GABRG2*), acetylcholine receptor signaling (*CHRM3, CHRNB4, CHRM1, CAMK2A*) and regulation of AMPA receptor activity (*GRIN3A, CACNG3, CACNG5, SHISA9*) (**Figure 5E**). In contrast, we observed downregulation of genes related to the synaptic vesicle cycle (*SLC1A6, SLC6A7, UNC13C, SLC6A3, CPLX1, CPLX3, CACNA1A, SLC18A2*), voltage gated potassium channels (*KCNA1, KCNA2, KCNA3*), glutamatergic receptors and regulators (*GRIA4, GRIN2C, GRIA2, GRM8, GRM4, CACNG4, CACNG7*) and serotonin receptors (*HTR3A, HTR5A*). Notably, patient prognosis appears to correlate with the expression of specific genes. Higher expression of *CHRNA1* was associated with poor prognosis, while lower expression of *CHRNA4, CHRNA6, GRIN2C, GABRE, GABRA1*, and *P2RX4* were linked to worst outcomes (**Figure 5K**). These findings suggest that communication with the neuronal microenvironment may play a beneficial role in this subgroup.

Group 4, the most common subgroup, showed less altered synaptic changes compared to other medulloblastoma subgroup (**Figure 5F**), with upregulation in synaptic vesicle cycle genes (*SLC32A1, SLC6A4, SLC17A6, SLC6A3, ATP6V0C, VAMP2, SLC6A5*), cholinergic signaling (*CHRM3, CHRM1, KCNQ5, CHRM2, CHRM4, CHRNA9*, *CAMK2A*), glutamatergic synapse (*SLC17A6, GRM7, GRM8, GRIN3A*) and neuropeptide Y signaling (*NPY2R, NPY5R*). In contrast, we observed downregulation of genes related to synaptic vesicle cycle (*SLC6A2, SYT1, CPLX1, SLC17A8, CPLX2, CPLX3, SLC18A1, SLC18A2*), GABAergic synapse (*GABRA4, ADCY8, HAP1, GAD2, GABRR1, KCNJ6, GABRG3*), dopaminergic synapse (*DRD2, DRD5*) and glutamatergic signaling (*GRIA4, ADCY8, SLC17A8, GRIN2A, GRIK3, GRIK2*). Interestingly, higher expression of specific synaptic receptors (*CHRNA5, GABRA5, GABRG2, GABRG3, GABRB3, GRIK1, NPY1R*) was associated with poor prognosis (**Figure 5H**), suggesting that targeting these pathways may represent a promising therapeutic strategy.

These findings reveal a complex, subgroup-specific rewiring of synaptic networks that underscores the crucial role of synaptic signaling in medulloblastoma biology and heterogeneity. Notably, the upregulation of **nAChR subunits** across all medulloblastoma subgroups and their association with poorer prognosis, mirroring patterns observed in gliomas, suggests that nAChR play a conserved role in tumor progression across diverse brain malignancies. In contrast, medulloblastomas exhibited more pronounced upregulation of muscarinic signaling, indicating distinct mechanisms of synaptic adaptation that may contribute to their unique clinical behavior. Given that cholinergic signaling influences both normal and malignant cell proliferation^39^, where muscarinic signaling enhances tumor growth and survival, while nicotinic signaling is more associated with differentiation, these findings underscore the potential for targeting cholinergic pathways in medulloblastoma treatment strategies. Further studies are needed to validate these associations and explore the therapeutic implications of targeting synaptic signaling in medulloblastoma treatment strategies.

### Ependymoma subtypes EPN-E1 and EPN-E2 exhibit divergent patterns of synaptic gene regulation with EPN-E1 showing a broad upregulation of genes related to synaptic vesicle cycling and GABAergic, glutamatergic, cholinergic, and NPY signaling

#### Subtype-specific enrichment of synaptic signaling in ependymoma subtypes

Ependymomas, arising from ependymal cells in the ventricular system and spinal cord, are not traditionally linked to neuronal circuit interactions. However, emerging evidence suggests they may influence nearby neuronal circuits ^17,40,41^. Symptoms such as seizures and cognitive deficits in patients with brain ependymomas ^42^, suggest potential neuronal disruptions via synaptic alterations or peritumoral effects.

Previous work has shown that using consensus clustering, the transcriptional data from ependymomas cluster into two primary compartments—EPN-E1 and EPN-E2^17^, where EPN-E1 is predominantly composed of supratentorial ependymomas and EPN-E2 is primarily dominated by posterior fossa ependymomas and less by NOS ependymoma. Following this compartmentalization in our study, EPN-E1 exhibited 68 upregulated and 22 downregulated synaptic genes (**Figure 6A, B, Supp Table 6**), while EPN-E2 displayed 22 upregulated and 61 downregulated synaptic genes compared to normal brain (**Figure 6C**). Interestingly, we identified 13 synaptic genes that were consistently upregulated across both EPN compartments (**Figure 6D**). Among these, we highlight *VAMP2* (**Figure 3F**), a key component of the SNARE complex, *CHRNA9* and *CHRM3* **(Figure 5J, K)**, both cholinergic receptors, and *SHISA9* and *CACNG3*, AMPA glutamate receptor complex members. Interestingly, *VAMP2*, *RPL36A*, *ATP6V0C*, *CHRNA9* and *HRNPH2* were also upregulated in glioma and medulloblastoma subgroups (**Figure 3F**, **Figure 5D, J, K**). In contrast, only one synaptic gene was consistently downregulated across the two EPN compartments, *CPLX3,* which is a complexin that regulates SNARE protein complex-mediated synaptic vesicle fusion (**Figure 6E**).

**Figure 6:**
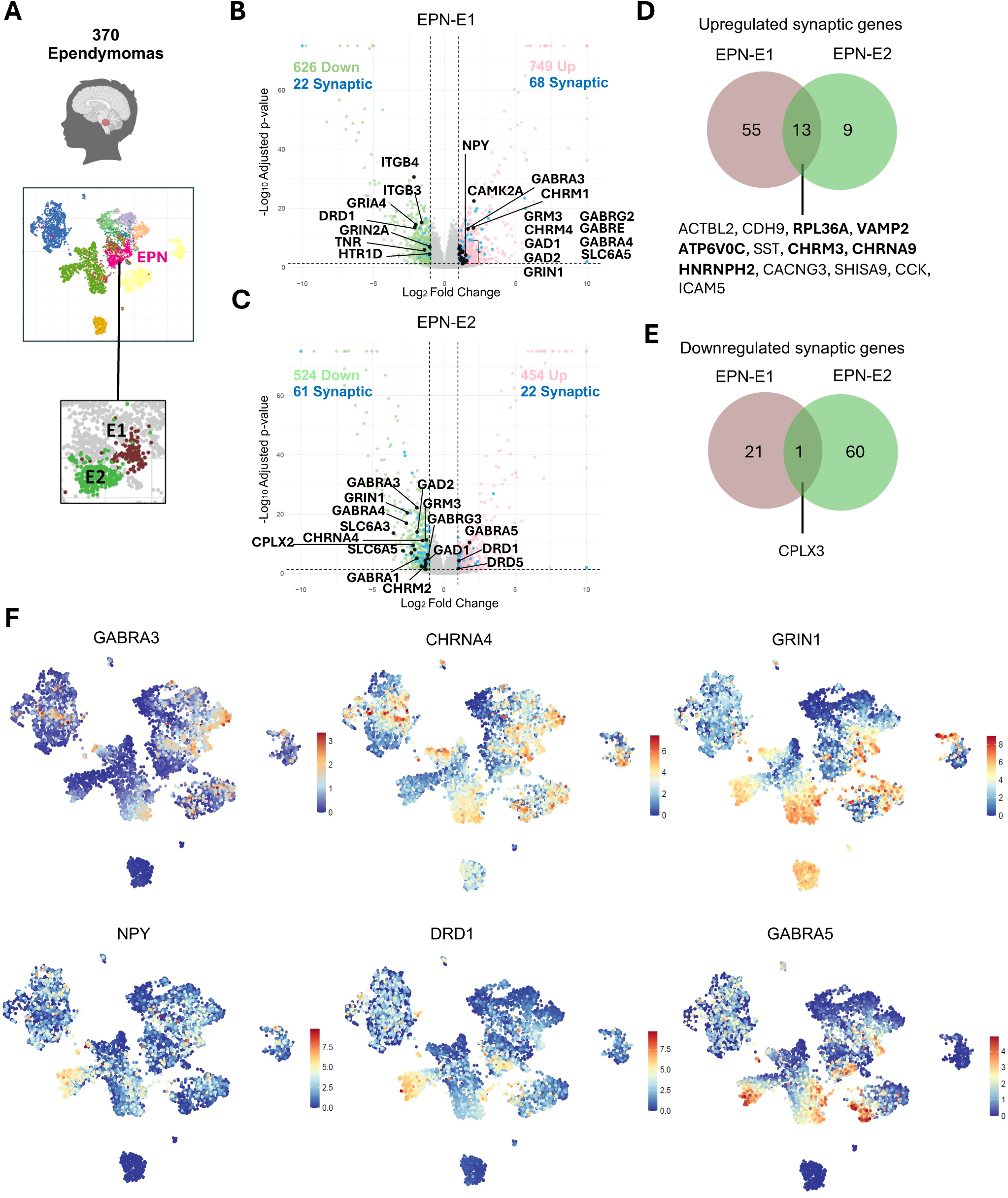
Distinct Synaptic Gene Expression Patterns Across Ependymoma Clusters. **(A)** Brain-UMAP representation highlighting ependymoma (EPN, pink) and two clusters (E1 and E2). **(B, C)** Volcano plots showing differentially expressed genes (DEGs) for ependymoma cluster: (**B**) EPN-E1 and (**C**) EPN-E2 compared to normal brain tissue (GTEx). Upregulated genes are shown in pink, downregulated genes in green, and differentially expressed synaptic genes are indicated in blue or black (highlighted). **(D, E)** Venn diagrams comparing shared (**D**) upregulated and (**E**) downregulated genes among the two ependymoma clusters. **(F)** Expression patterns of *GABRA3*, *CHRNA4*, *GRIN1, NPY, DRD1* and *GABRA5* visualized across the Brain-UMAP.

Focusing on EPN-E1 we observed an upregulation of synaptic genes related to GABA synthesis enzymes and receptor subunits (*GAD1, GAD2, GABRA3, GABRA4, GABRE, GABRG3*), cholinergic receptors (*CHRM1, CHRNA4, CHRM4*), glutamatergic receptors (*GRM3, GRIN1*), synaptic vesicle cycle (*SLC32A1, SLC6A2, SLC17A6, SLC6A3, SLC1A2, SLC18A3, SLC6A5*), synaptic plasticity (*CAMK2A*) and neuropeptide Y (*NPY*) (**Figure 6B**), which suggests a complex pattern of synaptic modulation. Intriguingly, NPY, a known modulator of neuronal activity, has been reported to suppress ependymoma tumor progression by remodeling the TME towards synaptic inhibition, thereby blunting brain hyperactivity and impeding tumor growth^40^. This interplay suggests that, while the upregulation of synaptic genes may indicate heightened neuronal signaling, NPY-mediated inhibitory effects might act as a counterbalance, promoting a less excitable TME. Notably, the downregulation of ionotropic glutamatergic receptors (*GRIA4, GRIN2A*), dopaminergic and serotoninergic receptors (*DRD1 and HTR1D)* and focal adhesion molecules (*ITGB4, TNR, ITGB3*) further supports a shift toward reduced excitatory signaling and diminished adhesion capacity.

In contrast, EPN-E2 exhibited a more limited impact on the upregulated synaptic gene network, characterized by the upregulation of dopaminergic receptors *(DRD1 and DRD5*) and a GABAergic receptor *GABRA5.* However, they showed an ample set of downregulated synaptic genes that included GABAergic synapses (*GABRA1, GABRA3, GABRA4, GABRG3, GAD1, GAD2*), glutamatergic receptors (*GRM3, GRIN1*), cholinergic receptors (*CHRNA4, CHRM2*) and synaptic vesicle cycle components (*SLC32A1, SLC6A2, SLC17A7, SLC6A7, SLC17A6, SLC6A3, CPLX2, SLC18A3, SLC6A5*) (**Figure 6C**), potentially reflecting a TME with diminished neural interactions and reduced synaptic functionality compared to EPN-E1.

Notably, most of the synaptic genes upregulated in EPN-E1 are downregulated in EPN-E2, suggesting that these subtypes exhibit fundamentally distinct synaptic landscapes (**Figure 6B, C, F**). These differences might be related to their anatomical location or differential interactions with the neural microenvironment. Although the small sample size precluded survival analysis, future studies investigating whether EPN-E1 is associated with poorer prognosis could yield valuable insights into its clinical relevance. These contrasting molecular profiles further highlight the heterogeneity within ependymomas and underscore the importance of gene expression-based subtype characterization to deepen our understanding of their biology and inform the development of targeted therapeutic strategies.

### Deregulation of synaptic genes in meningiomas indicates a heightened synaptic connectivity and remodeling, with lower expression linked to better prognosis

Although meningiomas are extra-axial tumors, their proximity to the brain parenchyma can indirectly impact neuronal function and synaptic plasticity. They often compress adjacent brain tissue and cause peritumoral edema, exacerbating neuronal damage and potentially influencing tumor progression^43^.

In our DEG analysis, we observed a significant upregulation of several glutamate receptor genes, including *GRID2*, *GRIK3*, and *GRIA4*, in meningiomas compared to normal brain tissue (**Figure 7B, Supp Table 7**). These genes encode delta, kainate and AMPA receptor subunits, which are critical mediators of fast excitatory synaptic transmission. Their upregulation suggests heightened excitatory neurotransmission, potentially contributing to increased neuronal excitability and susceptibility to seizures^44^. In contrast, we identified downregulation of *GRIN1* (**Figure 6F**), *GRIN2A*, and *GRIN2B* (**Figure 7B**), which encode subunits of NMDA receptors that are crucial for synaptic plasticity. While kainate and AMPA receptors mediate rapid excitatory currents, NMDA receptors are slower and more modulatory^45^. Their suppression may result in an inability to effectively regulate or terminate excitatory inputs, further exacerbating neuronal excitability.

**Figure 7:**
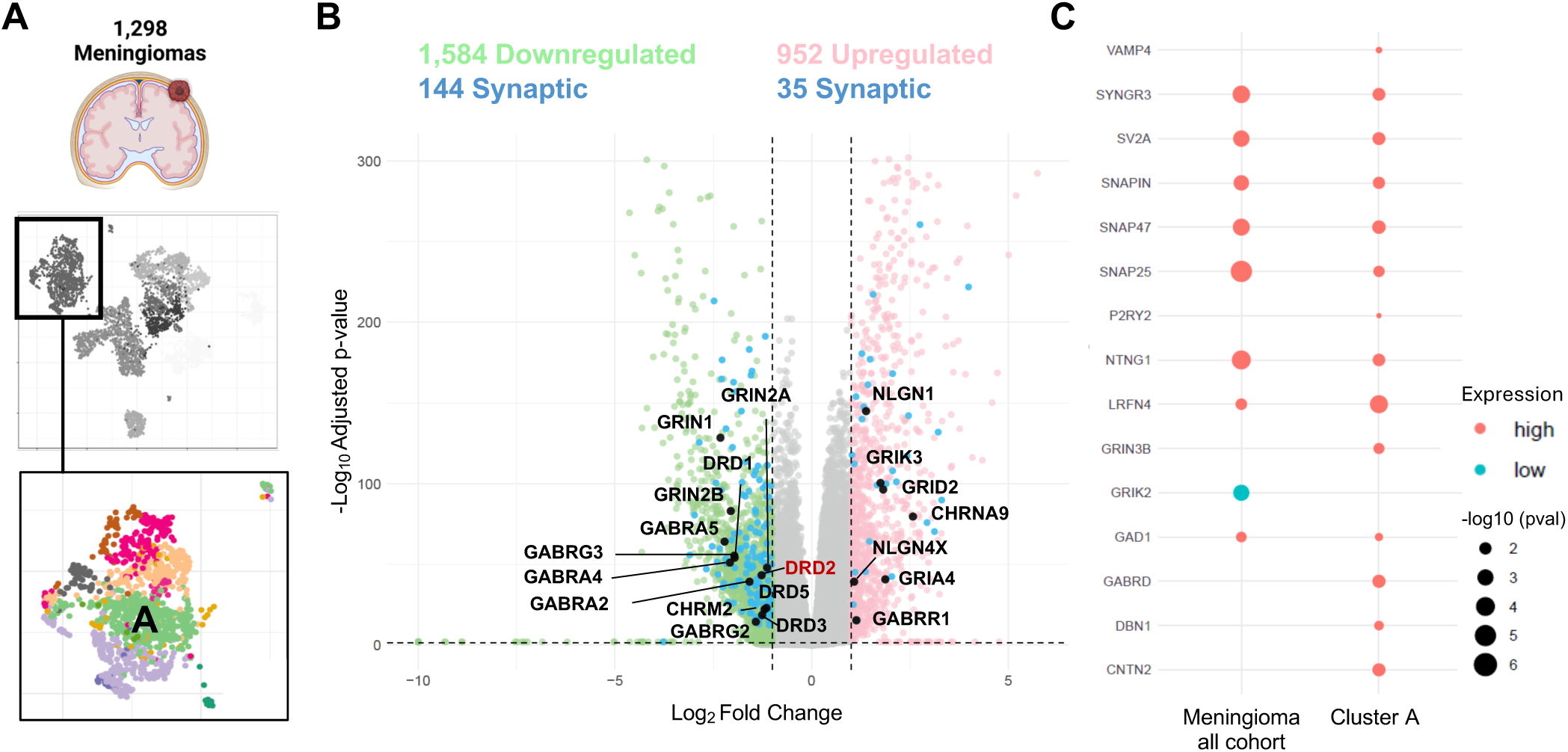
Synaptic Gene Expression Patterns and Clinical Associations in Meningiomas. **(A)** Brain-UMAP representation highlighting meningiomas (square) and their respective clusters shown in distinct colors, including Cluster A, which is associated with poorer survival. **(B)** Volcano plot showing differentially expressed genes (DEGs) for meningioma compared to normal brain tissue (GTEx). Upregulated genes are shown in pink, downregulated genes in green, and differentially expressed synaptic genes are indicated in blue or black (highlighted). **(C)** Dot plot showing the association between synaptic receptor gene expression and survival in the overall meningioma cohort and in cluster A. Only synaptic genes with statistically significant p-values are displayed (*p* < 0.05). Genes whose higher expression is associated with poorer survival are shown in orange, while those linked to better prognosis are shown in turquoise. Dot size represents the statistical significance of the association, expressed as –log₁₀(p-value).

Alterations in GABAergic signaling were also identified. Specifically, we found upregulation of *GABRR1* and *GABRA6*, which are associated with inhibitory synaptic transmission. However, several more GABA receptor subunit genes such as *GABRG1*, *GABRG2*, *GABRG3, GABRA2*, *GABRA4*, *GABRA5* (**Figure 6F**) and *GABRQ* were markedly downregulated (**Figure 7B**). This dysregulation could lead to ineffective or inconsistent inhibitory tone, contributing to the overall imbalance between excitatory and inhibitory signaling.

In terms of cholinergic neurotransmission, we observed an upregulation of *CHRNA9*, a receptor that facilitates cation influx and excitatory signaling and is also upregulated in other brain tumor types (**Figure 5J**). Conversely, we found a downregulation of *CHRM2* and *CHRM5*, both of which regulate neuronal excitability (**Figure 7B**). The reduced expression of *CHRM2* could diminish inhibitory signaling, while the downregulation of *CHRM5* may impair excitatory modulation, exacerbating the excitatory-inhibitory imbalance and potentially promoting neuronal hyperexcitability.

Interestingly, we also observed an upregulation of neuroligins *NLGN4X*, *NLGN44* and *NLGN1* (**Figure 7B**), which are essential synaptic cell-adhesion molecules that mediate the formation, maturation, and stabilization of synapses. It is also important to note that neuroligin imbalance has been implicated in epilepsy^46,47^ and gliomas ^2,4^. Their upregulation could suggest an environment favoring heightened synaptic connectivity or remodeling, potentially as a compensatory mechanism in response to neuronal compression or edema.

Additionally, we observed a downregulation of genes related to the dopaminergic system, including *DRD1*, *DRD2*, *DRD3*, and *DRD5* (**Figure 7B**). Given that the meninges lack significant dopaminergic innervation, the baseline expression of dopamine receptors in meningeal cells or meningiomas may inherently be low, potentially accounting for the observed differences when compared to brain parenchyma. In this sense, since meningiomas arise from arachnoid cap cells of the arachnoid mater, directly comparing them to normal brain tissue may not provide the most meaningful insights, representing a limitation of the study. However, when we evaluate the influence of synaptic gene expression on survival outcomes, our analysis revealed that lower expression of key SNARE (*SNAP25, SNAP47, SNAPIN*), synaptic vesicle-associated (*SV2A, SYNGR3),* neurite outgrowth (*NTNG1, LRFN4*), GABA-related genes (*GAD1)* correlated with improved patient survival, whereas high *GRIK2*, expression was associated with better prognosis (**Figure 7C**). Given the association of cluster A meningiomas with poorer prognosis^19^, we examined the expression of synaptic receptors in this cluster compared to the overall meningioma cohort (**Figure 7C**). While general trends remained consistent, the distinct prognostic impact of *VAMP4*, *DBN1*, *CNTN2*, *GRIN3B*, *GABRD* and *P2RY2* in Cluster A, where higher expression correlated with poor prognosis, suggests that synaptic gene expression may play a pivotal role in shaping meningioma progression (**Figure 7D**).

### Single-Cell Data Suggest Tumor Cell Expression of Synaptic Receptor Genes

One of the main challenges when interpreting bulk RNA-seq data from CNS tumors is determining whether the detected signal originates from tumor cells themselves or from neighboring neuronal populations that may innervate the tumor. To address this, we selected synaptic receptor genes that were differentially expressed across glioma and medulloblastoma subgroups and examined their mean expression in independent single-cell RNA-seq datasets, in which expression profiles predominantly represent tumor cells. For example, *GRIA4*, a postsynaptic AMPA receptor subunit, is downregulated in GBM, WNT, and Groups 3 and 4 medulloblastomas but upregulated in SHH tumors in bulk RNA-seq. Single-cell data corroborate this pattern, showing that a substantial fraction of cells in SHH medulloblastomas express *GRIA4* at high levels **(Supplementary Figure 8).** Similarly, *GRIK2*, a kainate-type glutamate receptor, *GRIN2A*, a postsynaptic NMDA receptor subunit, and *CHRNA5*, a postsynaptic nicotinic cholinergic receptor subunit, are expressed at high levels in a large proportion of SHH cells, mirroring the bulk RNA-seq findings.*CHRNA9*, another postsynaptic acetylcholine receptor subunit, is markedly upregulated in GBM, WNT, and Group 4 medulloblastomas but downregulated in low-grade gliomas. Single-cell analyses reveal that only a subset of cells express *CHRNA9*, yet those that do show high expression levels, suggesting context-dependent regulation or partial neuronal contribution. Finally, *HNRNPH2*, which regulates RNA processing and synaptic gene expression, is broadly upregulated across GBM, LGG, and all medulloblastoma subgroups, with a high proportion of cells expressing it in single-cell datasets, albeit with variable intensity. Together, these consistent expression patterns across bulk and single-cell datasets indicate that the synaptic receptor signatures detected in CNS tumors are predominantly tumor cell–derived, although contributions from infiltrating or adjacent neurons cannot be fully excluded, as observed for *CHRNA9*.

## Discussion

The role of synaptic interactions between tumor cells and the TME has long been underestimated. However, the emergence of the cancer neuroscience field, driven by elegant and cutting-edge studies in the field, is reshaping this perspective, revealing the profound impact of neuronal activity on tumor progression and resistance mechanisms ^1–5,7,8,10,11,13,16^. Our study presents, for the first time, a comprehensive transcriptomic analysis of 4,330 adult and pediatric brain tumor samples and 1,409 brain samples, demonstrating how different brain tumor types regulate synaptic pathways within the TME, highlighting both shared and tumor-specific mechanisms. This reveals a complex landscape where some tumors predominantly upregulate synaptic genes, others exhibit widespread downregulation, and some display both patterns. The functional and physiological implications of these alterations remain to be elucidated via mechanistic studies. Increased synaptic signaling of specific receptors may suggest enhanced tumor-neuron interactions that support tumor progression, whereas the downregulation of other synaptic receptors could indicate disrupted neuronal connectivity or reduced synaptic input within the TME. Alternatively, the downregulation of synaptic genes, particularly adhesion molecules, may reflect weakened cell-cell or synaptic interactions, potentially affecting tumor migration or invasion. It could also suggest a developmental arrest or a reversion to a less differentiated, more progenitor-like state, a hallmark of aggressive cancer subtypes. These findings highlight the need for further functional studies to elucidate the precise role of synaptic remodeling in brain tumor biology.

In recent years, RNA-seq has become a fundamental tool for profiling gene expression across various diseases, including cancer^17–19^. While scRNA-seq offers critical insights into the cellular heterogeneity within tissues, its application is often limited by high costs, extensive bioinformatic work, and small sample sizes. In contrast, leveraging existing bulk RNA-seq datasets to construct reference landscapes offers a more cost-effective and scalable alternative, enabling the study of gene expression patterns across substantially larger cohorts. This approach has allowed us to systematically explore alterations in synaptic gene expression and signaling networks, shedding light on their role in tumor subtype progression and unveiling new opportunities for biomarker discovery and targeted therapy development. Additionally, this RNA-seq dataset will be publicly available through Oncoscape, offering a valuable resource for research groups with limited access to or expertise in bioinformatics tools. This accessibility could enhance the impact of numerous studies accelerating scientific progress.

Our findings reveal that *HNRNPH2* is consistently upregulated across gliomas and medulloblastomas, pointing to a possible role in brain tumor progression. Mutations in *HNRNPH2* cause an X-linked neurodevelopmental disorder characterized by intellectual disability, autism-like behaviors, and epilepsy, all conditions involving synaptic dysfunction^48,49^. Like other heterogeneous nuclear ribonucleoproteins (hnRNPs), *HNRNPH2* regulates alternative splicing of transcripts involved in neuronal function^34^, as well as dendritic arborization^49^. hnRNPs have been shown to modulate the splicing of genes like *GRIN1* (encoding an NMDA receptor subunit) and *CAMK2* (calcium/calmodulin-dependent protein kinase II)^50,51^, both essential for synaptic signaling and plasticity. Additionally, *HNRNPH2* plays a role in mRNA stability and transport^34,48^, processes crucial for localized synaptic protein synthesis. Given its function in RNA processing^34,48^, *HNRNPH2* may drive tumor-specific alternative splicing programs that enhance malignancy by promoting proliferation, invasion, and therapy resistance. Furthermore, it may dysregulate the expression of synaptic receptors, adhesion molecules, or ion channels, altering the TME and facilitating tumor progression. Targeting expression of mutant *HNRNPH2*, such as with an antisense oligonucleotide, has shown promising results in mitigating the gain-of-function effects associated with certain *HNRNPH2* mutations linked to neurodevelopmental disorders^49,52^. Future translational studies should explore the therapeutic potential of therapeutic approaches that aim to reduce the expression of *HNRNPH2* or inhibit its functional signaling in brain tumors.

In our study on synaptic signaling adaptations across glioma subtypes, we show that P2 purinergic receptor subtypes, key regulators of synaptic function, are consistently downregulated in both adult and pediatric gliomas compared to normal brain tissue. Notably, high expression of P2X receptor subtypes correlates with poorer prognosis, suggesting a potential role in tumor progression. Beyond gliomas, P2 purinergic receptors have been implicated in cancer progression, influencing ATP-driven proliferation, immune evasion, inflammation ^53,54^. P2XRs are typical ligand ATP-gated cation channels allowing Na^+^, K^+^ and Ca^2+^ fluxes across the plasma membrane ^55^. There is growing evidence that P2 purinergic receptor expression within the TME is crucial for both disease initiation and progression, as multiple immune cells within the TME express P2 receptors, influencing tumor-immune interactions ^55^. Consequently, several clinical trials are currently investigating purinergic signaling inhibitors as stand-alone or combination cancer therapies ^54^. However, the results have been largely disappointing ^55^, likely due to the widespread distribution of P1 and P2 receptors across cancer and host immune cells, leading to unintended effects. A deeper understanding of this complex and understudied tumor-neuron-immune signaling network will be essential for the design of next-generation purinergic-targeted therapies to reprogram the TME and improve clinical outcomes.

Our findings reveal that nAChR dysregulation is associated with glioma aggressiveness in both adult and pediatric patients. Among these receptors, *CHRNA9* emerges as a key player, showing significant upregulation not only in glioma progression but also in WNT and Group 4 medulloblastomas, ependymomas, and meningiomas. This widespread involvement across multiple brain tumor types highlights *CHRNA9* as a promising therapeutic target for disrupting tumor-promoting pathways and restoring a more homeostatic TME. While *CHRNA9* has been extensively studied in other malignancies, including melanoma, breast and lung cancers^56^, its role in brain tumors remains poorly understood. In non-brain cancers, *CHRNA9* activation has been linked to pro-tumorigenic effects, such as enhanced proliferation, invasion, and resistance to apoptosis^56^. Recently, *CHRNA9* has been shown to be upregulated in glioma tissues, where its expression correlates with poor prognosis and reduced patient survival^57^. Here, we provide evidence that *CHRNA9* is not only relevant in gliomas but also plays a role in other brain tumor types, underscoring its potential as a novel target for therapeutic intervention across multiple brain malignancies. While *CHRNA9* receptors were found in the hair cells of the inner ear and regulate auditory functions^58^, their expression was later observed in many other locations and tissues^56,59^. CHRNA9 encodes a functional ion channel that mediates Ca²⁺ influx, influencing synaptic plasticity and signal transduction^60^. Unlike most neuronal nAChRs, *CHRNA9* does not form classical heteromeric receptors in the brain but instead heteromerizes with *CHRNA10*, conferring unique pharmacological properties^58^. *CHRNA9* is also expressed in immune cells^61^ and emerging evidence suggests that *CHRNA9* plays a role in exacerbating disease in inflammatory and autoimmune responses^62^, which may critically influence tumor-neuron interactions. Given these findings, α9-nAChR antagonists are being explored in cancer therapy for their potential to inhibit tumor-promoting signaling^56^. Targeting *CHRNA9* in brain tumors could represent a novel therapeutic strategy, potentially in combination with existing treatments. However, further research is needed to fully elucidate its role and therapeutic potential in brain tumors. In summary, our findings highlight the complexity of neuron-tumor crosstalk and its potential influence on patient survival and brain tumor plasticity. By identifying key synaptic signaling pathways altered across tumor types, such as *HNRNPH2,* P2 purinergic signaling and cholinergic signaling including *CHRNA9*, this study lays the foundation for future investigations into how targeting these pathways could lead to novel therapeutic strategies. A promising avenue for the field is leveraging existing therapies developed for neurological disorders and combining them with established treatments that target tumor-intrinsic signaling mechanisms^11,12^. Further research will be critical to determining whether modulating specific synaptic components can reprogram the TME toward an anti-tumoral state, ultimately improving patient outcomes. To fully explore the findings presented here, there is a need for enhanced collaborations and crosstalk between synaptic biologists and scientists with expertise in brain tumors and cancer, fostering a multidisciplinary approach to uncover novel therapeutic strategies.

## Acknowledgements

We thank members of the Pattwell Lab at Seattle Children’s Research Institute (SCRI) and the Holland lab at Fred Hutchinson Cancer Research Center (FHCRC) for valuable discussions, and collaborators for sharing their data and metadata. This research was supported by funding from the FHCC (E.C.H) and Seattle Children’s Research Institute (S.S.P., L.M.G.) and University of Washington School of Medicine (S.S.P., L.M.G.).

## Author Contributions

Conceptualization: L.M.G., S.S.P.; Methodology: L.M.G, S.A., G.G., M.J., E.C.H., S.S.P. Investigation: L.M.G, S.A.; Bioinformatic analysis: S.A.; Visualization: L.M.G., S.A. G.G., M.J.; Funding acquisition: E.C.H., S.S.P.; Project administration: L.M.G., S.A.; Supervision: S.S.P.; Writing – original draft: L.M.G., S.A.; Writing – review & editing: L.M.G., S.A., G.G., M.J., E.C.H., S.S.P.

## Data and materials availability

All analysis including statistics and visualization were done in R version 4.3. Plots were generated using R basic graphics and ggplot2. All custom code used in this study are available at https://github.com/sonali-bioc/ComprehensiveBrainUMAP

---

**Supplemental Figure 1:**
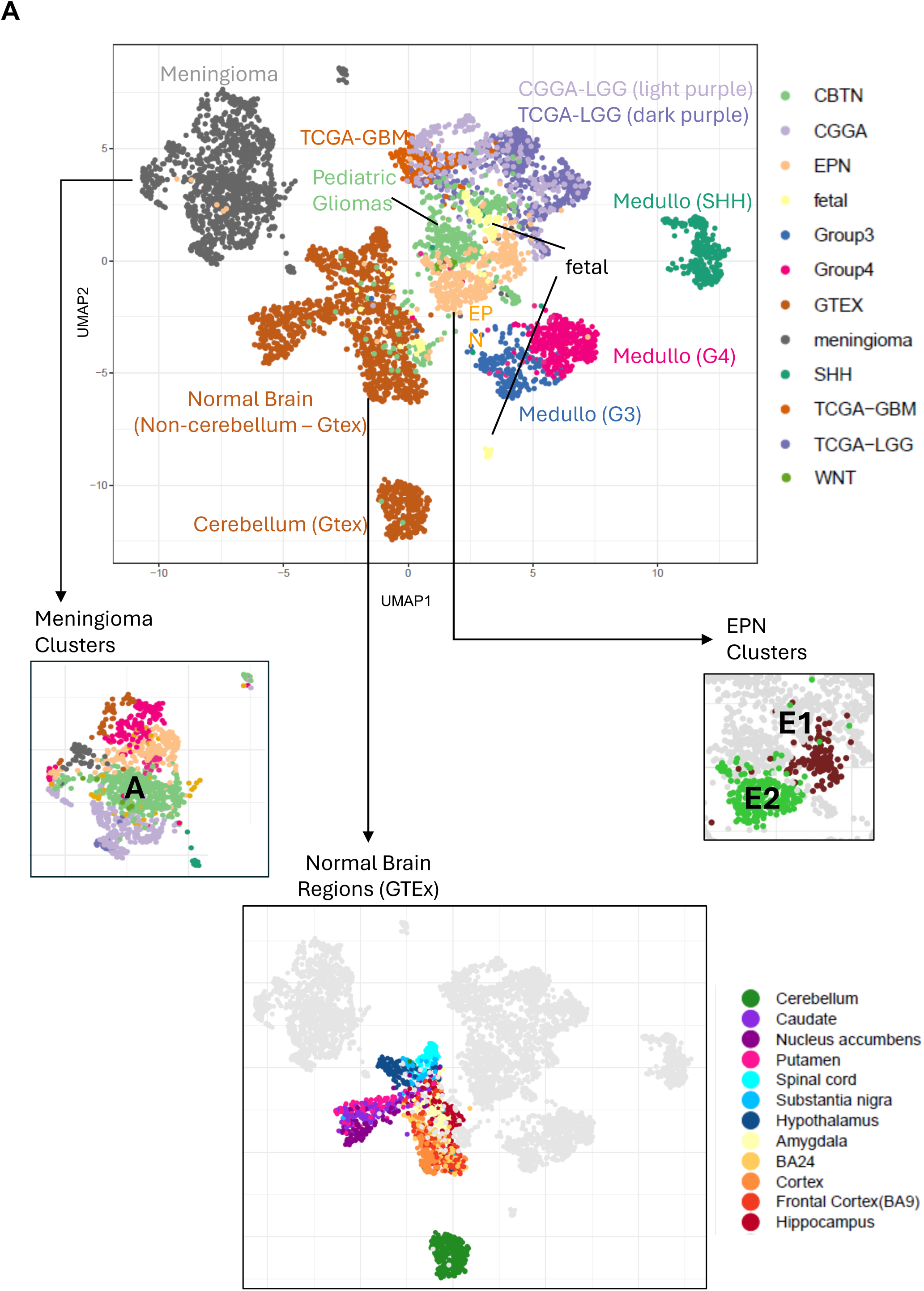
UMAP Representation of the Full Dataset and Subclusters. **(A)** Brain-UMAP representation of the complete dataset, including adult gliomas from TCGA-LGG (The Cancer Genome Atlas Low Grade Glioma), TCGA-GBM, and CGGA-LGG (Chinese Glioma Genome Atlas); pediatric gliomas; medulloblastoma subgroups (SHH, Group 3, Group 4, and WNT); ependymoma subclusters (EPN-E1 and EPN-E2); and meningioma clusters, including Cluster A, which is associated with poorer survival. Normal brain regions from GTEx (Genotype-Tissue Expression project) are also included. EPN, ependymoma; GBM, glioblastoma; GTEx, Genotype-Tissue Expression project; LGG, low-grade glioma; SHH, sonic hedgehog; TCGA, The Cancer Genome Atlas Low Grade Glioma.

**Supplemental Figure 2:**
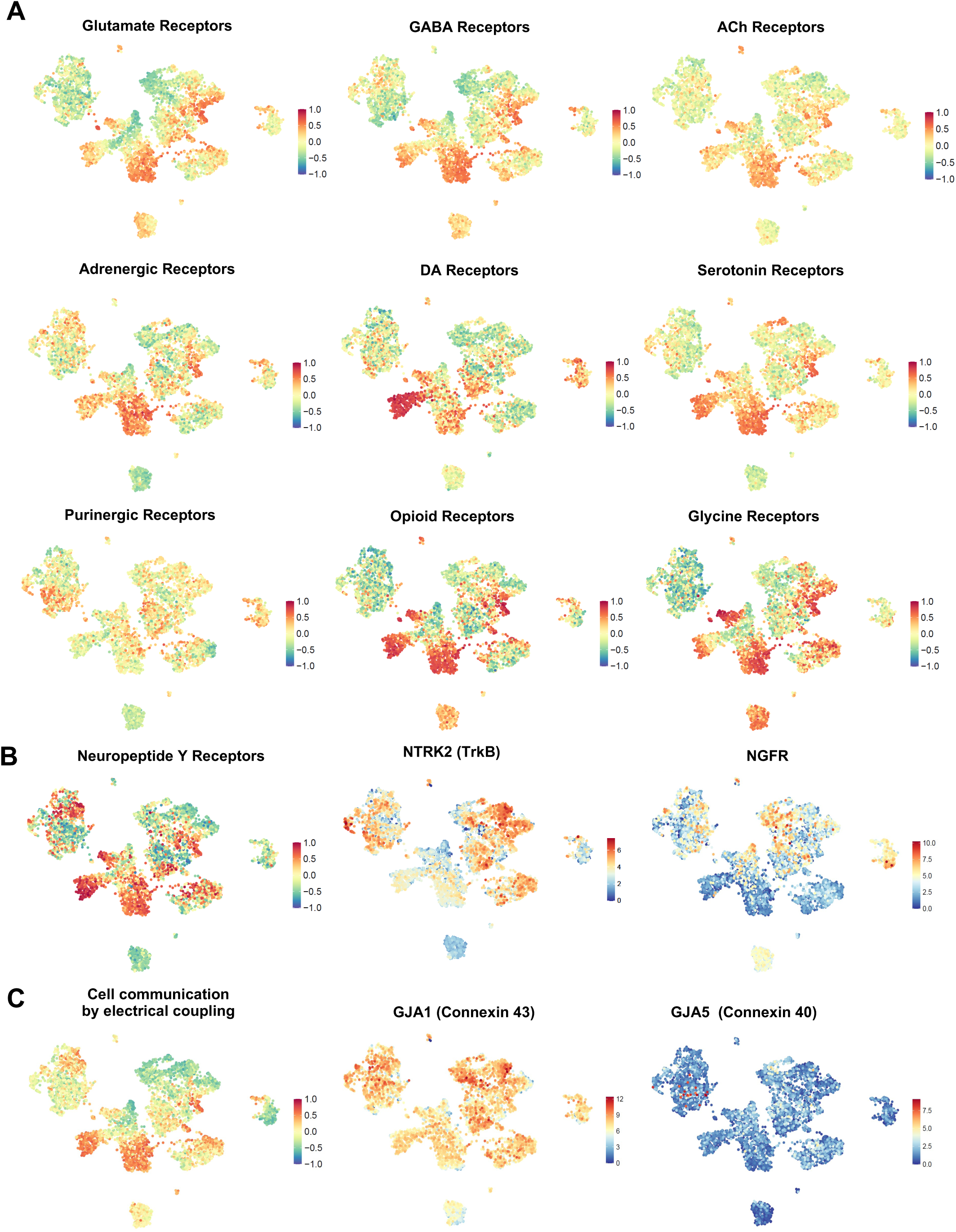
Neuronal Signatures in Brain Tumors. **(A, B,C**) Gene Set Variation Analysis (GSVA) visualized in the Brain-UMAP using selected gene sets from Harmonizome 3.0, including: (**A**) neuronal receptor-related genes including glutamate, GABA, acetylcholine (ACh), adrenergic, dopamine (DA), serotonin, purinergic, opioid and glycine receptors, (**B**) paracrine signaling genes involving Neuropeptide Y (NPY) receptors, *NTRK2* and *NGFR* genes, and (**C**) genes related to electrical coupling, and specifically *GJA1* and *GJA5*. GSVA scores range from −1 to 1 per sample.

**Supplemental Figure 3:**
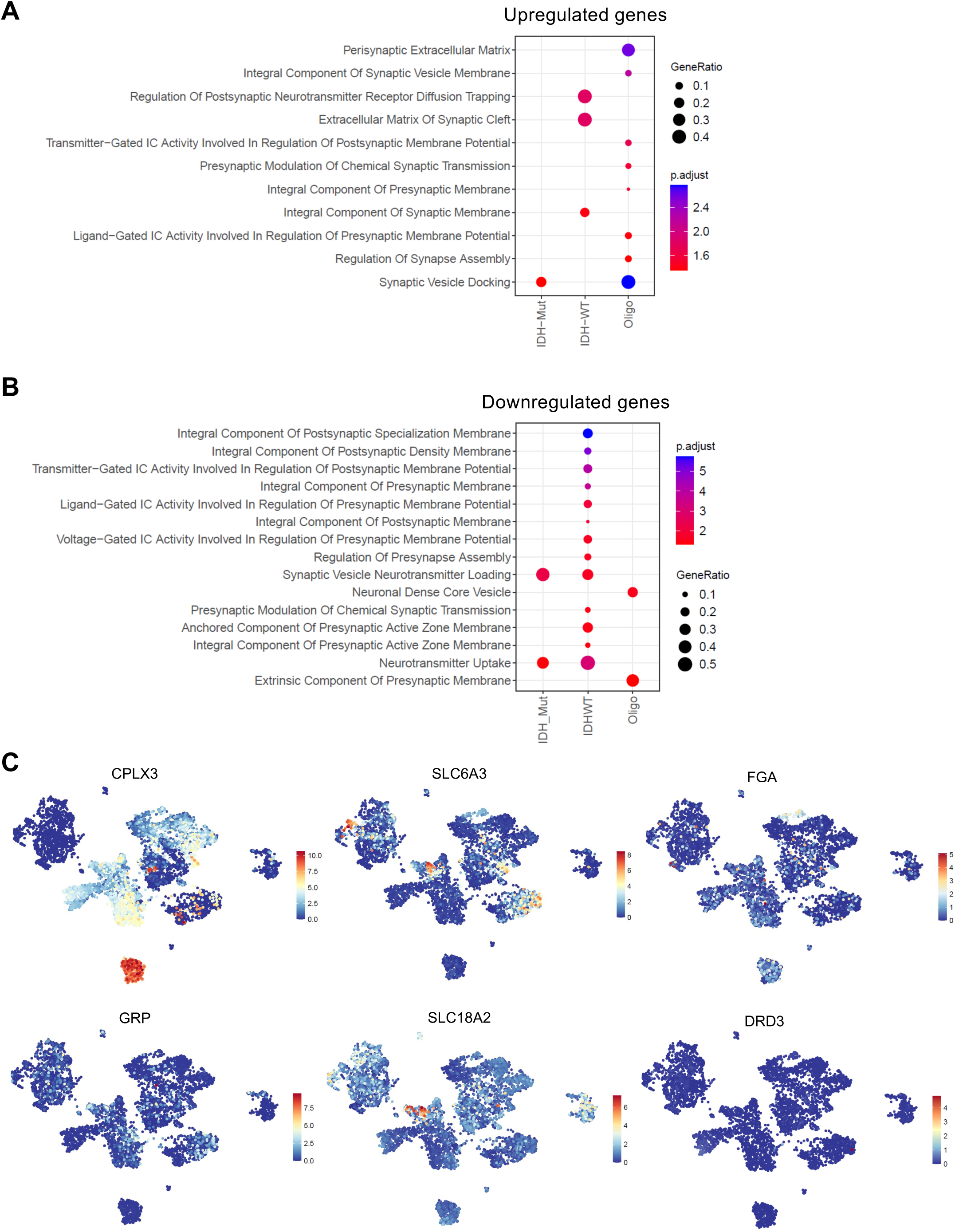
Synaptic Gene Ontology Analysis of Upregulated and Downregulated genes in Adult Gliomas. **(A, B)** Synaptic Gene Ontology (SynGO) analysis for upregulated (A) and downregulated (B) genes in astrocytoma, GBM and oligodendroglioma. **(C)** Expression patterns of shared downregulated synaptic genes (*CPLX3, SLC6A3, FGA, GRP, SLC18A2, DRD3*) for the three adult glioma types visualized across the Brain-UMAP.

**Supplemental Figure 4:**
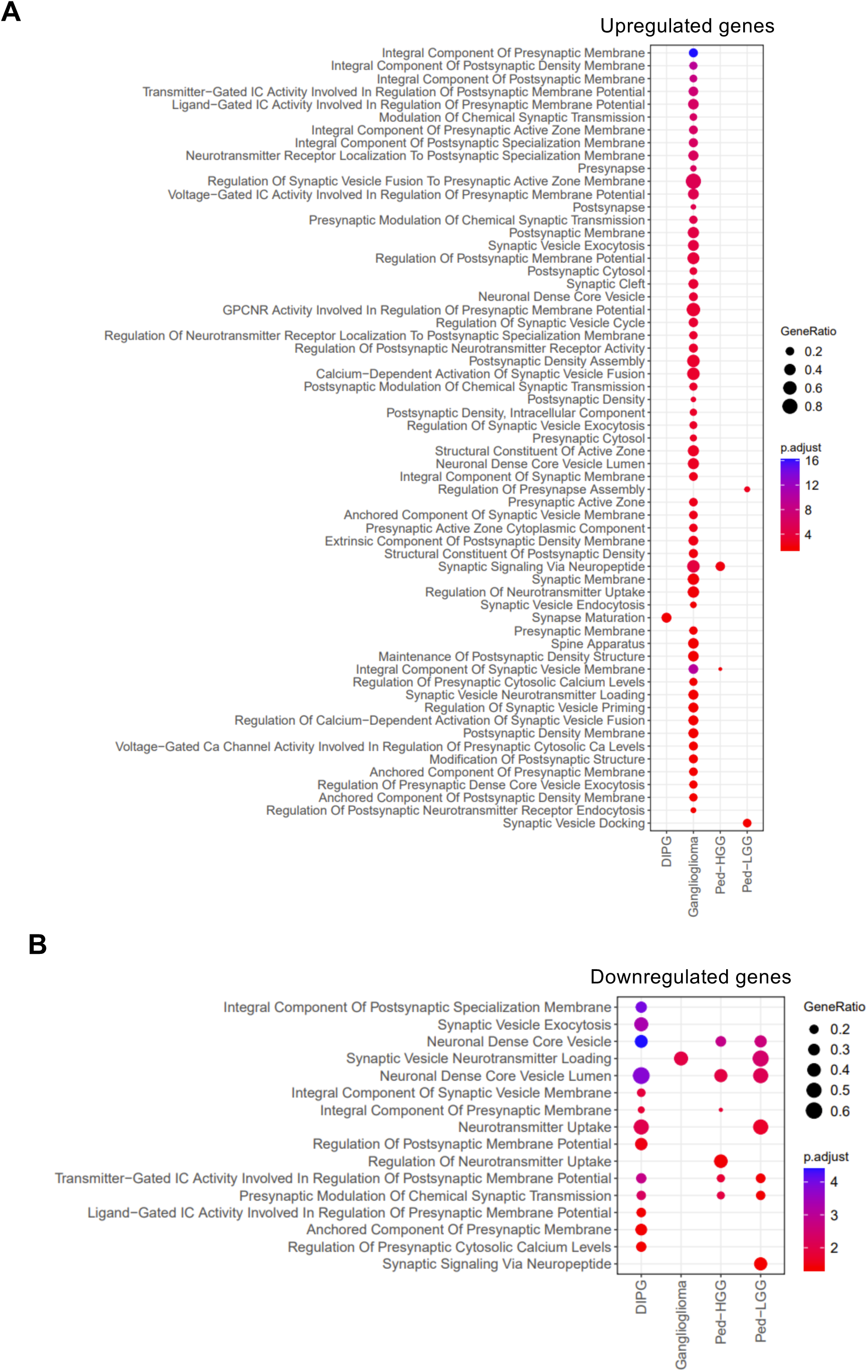
Synaptic Gene Ontology Analysis of Upregulated and Downregulated genes in Pediatric Gliomas. **(A, B)** Synaptic Gene Ontology (SynGO) analysis for upregulated (**A**) and downregulated (**B**) genes in DIPG, ganglioglioma, pediatric high-grade glioma (P-HGG) and pediatric low-grade glioma (P-LGG).

**Supplemental Figure 5:**
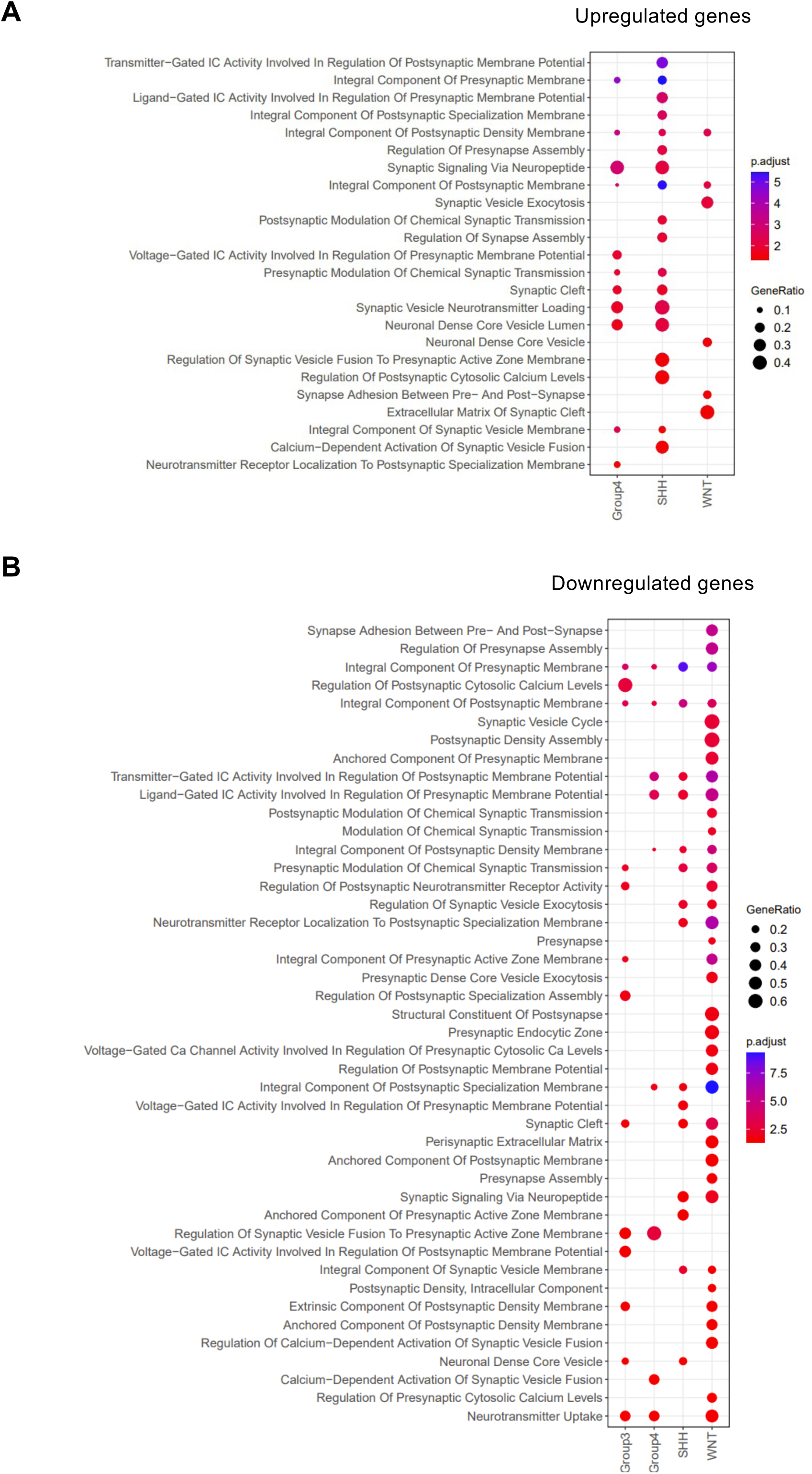
Synaptic Gene Ontology Analysis of Upregulated and Downregulated genes in Medulloblastoma. **(A, B)** Synaptic Gene Ontology (SynGO) analysis for upregulated (**A**) and downregulated (**B**) genes in medulloblastoma subgroups: Group 3, Group 4, SHH and WNT.

**Supplemental Figure 6:**
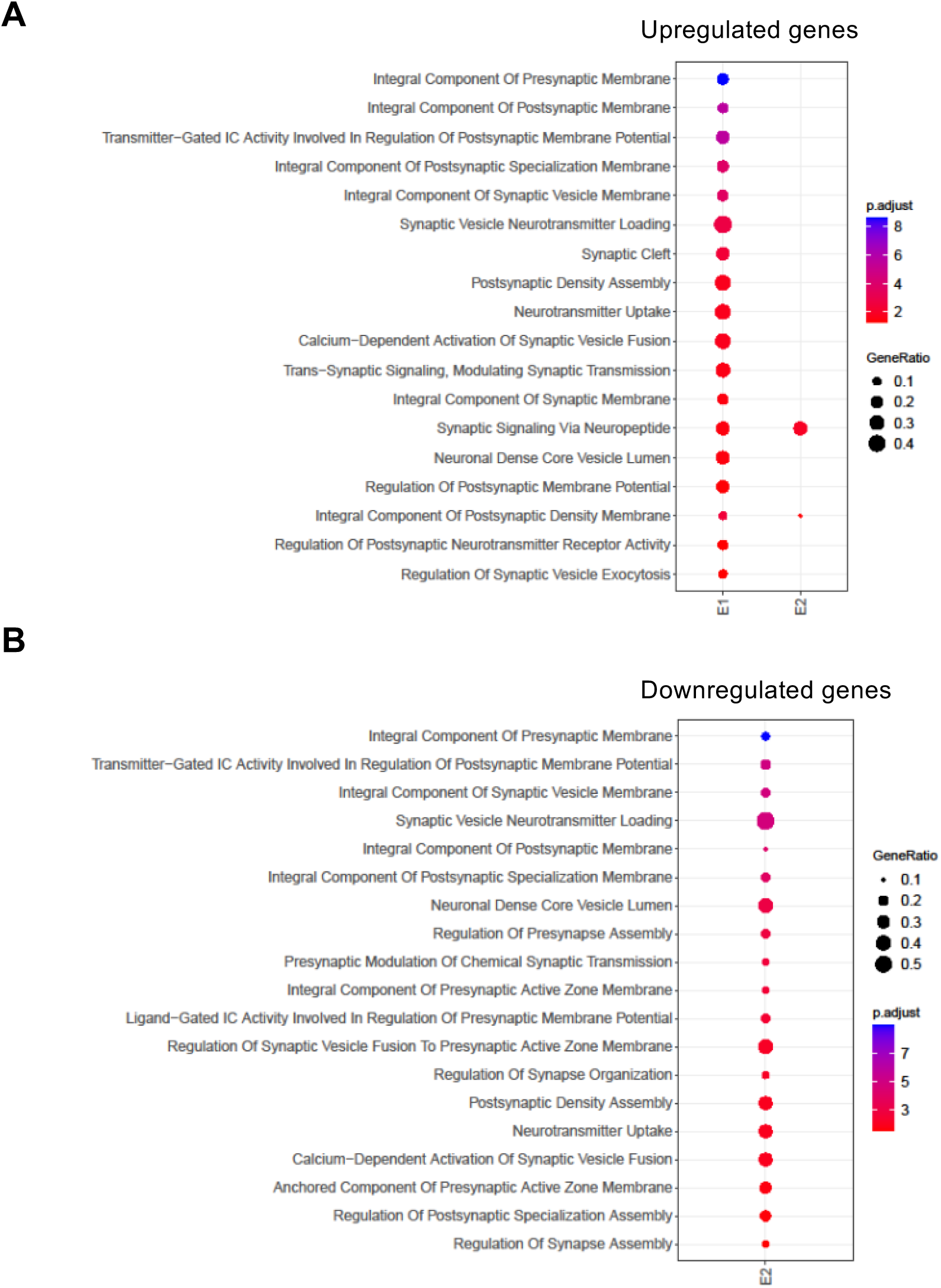
Synaptic Gene Ontology Analysis of Upregulated and Downregulated genes in Ependymoma. **(A, B)** Synaptic Gene Ontology (SynGO) analysis for upregulated (**A**) and downregulated (**B**) genes in ependymoma clusters: E1 and E2.

**Supplemental Figure 7:**
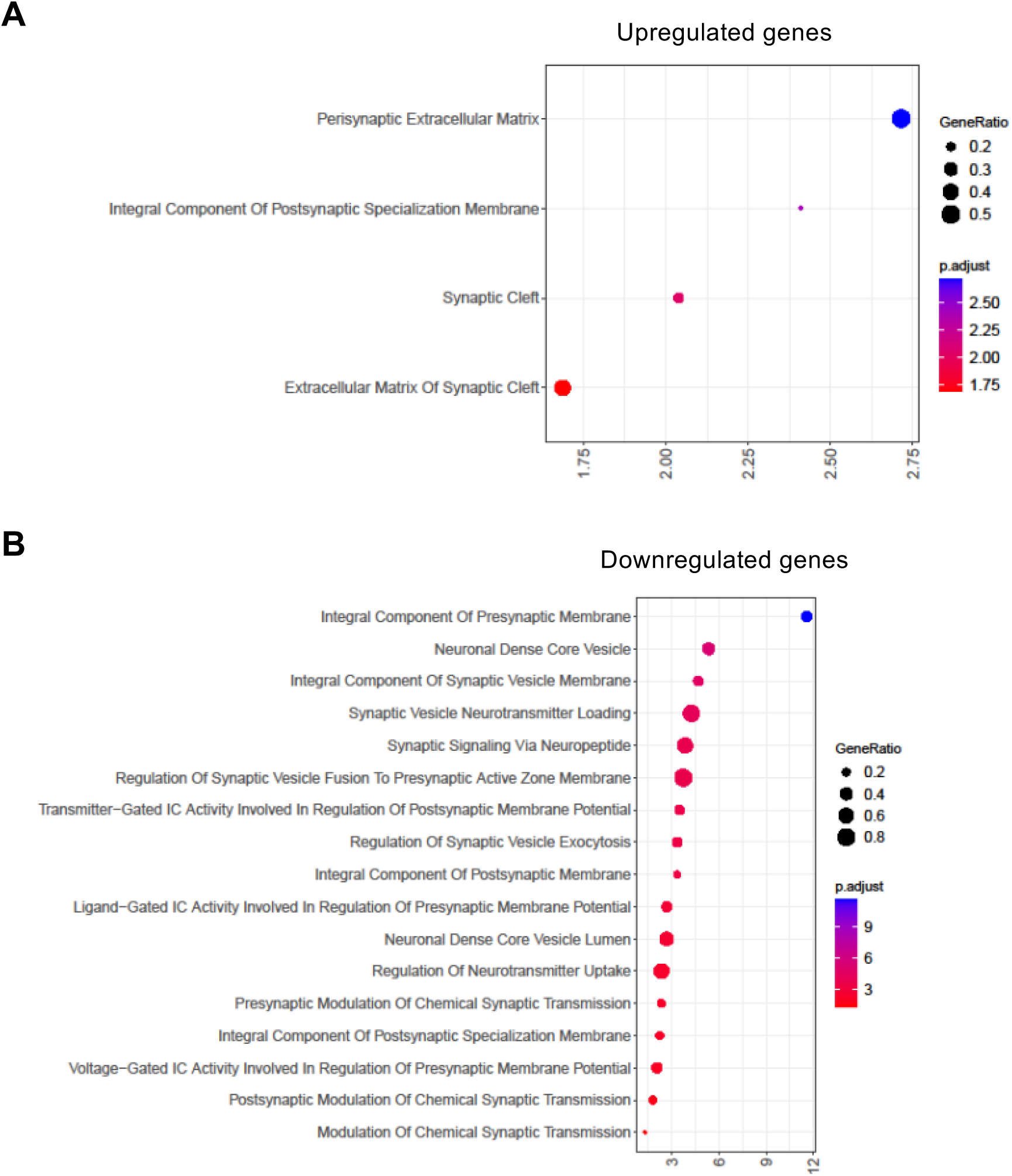
Synaptic Gene Ontology Analysis of Upregulated and Downregulated genes in Meningioma. **(A, B)** Synaptic Gene Ontology (SynGO) analysis for upregulated (**A**) and downregulated (**B**) genes in meningioma.

**Supplemental Figure 8:**
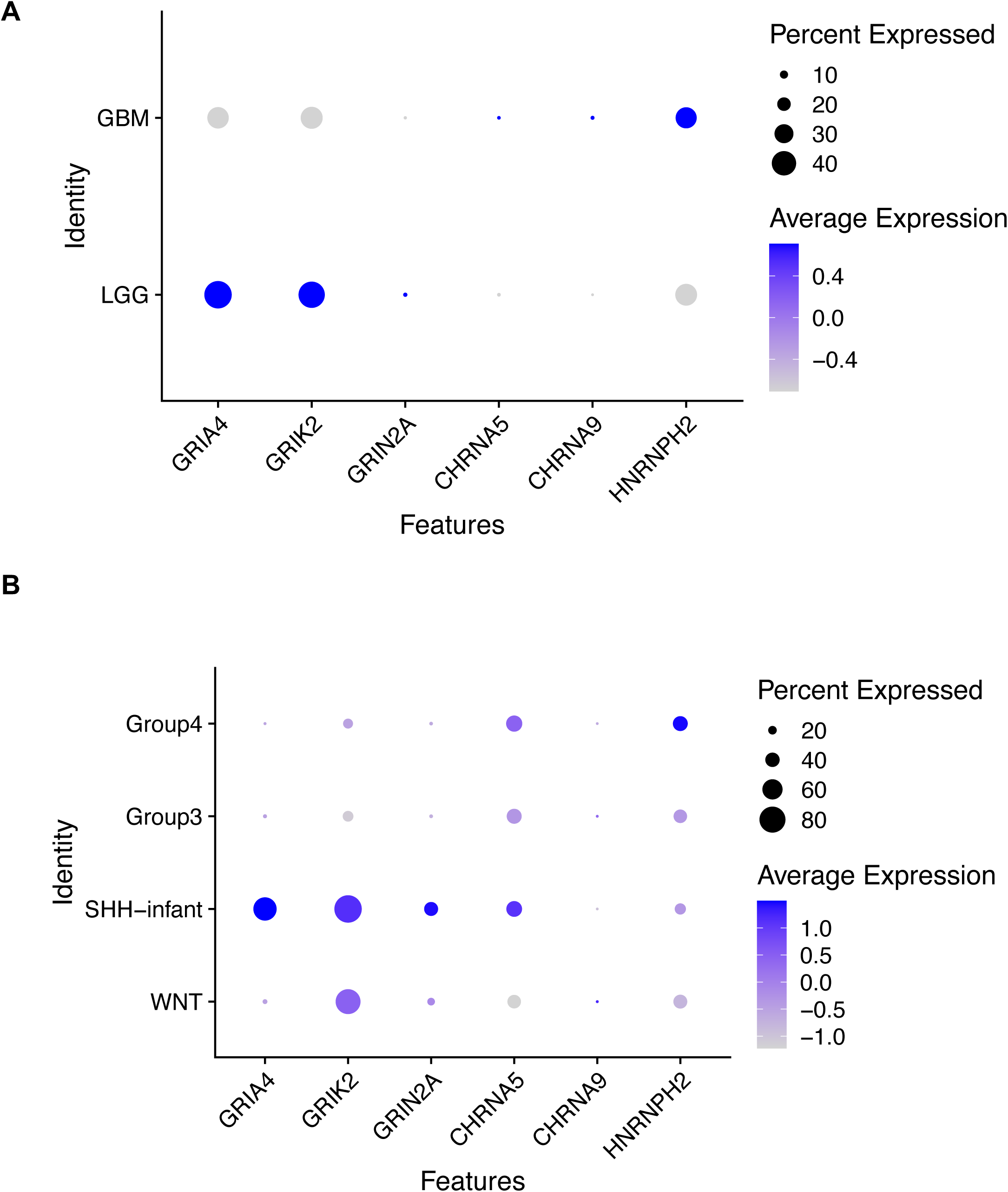
Mean single-cell expression of synaptic genes. (**A**) Dot plot showing the mean expression of selected synaptic genes in adult gliomas (GBM and LGG). (**B**) Dot plot showing the mean expression of selected synaptic genes across medulloblastoma subgroups.

## Table Legends

**Supplemental Table 1. Distribution of Tumor Samples Across Histopathological Categories**

*Summary of the number of tumor samples classified by histopathological subtype*.

**Supplemental Table 2. Catalogue of Synaptic Genes and Receptor Targets Analyzed in This Study**

*Comprehensive list of synaptic genes and receptor families included in the transcriptomic analyses.*

**Supplemental Table 3. Differential Gene Expression Profiles in Adult Glioma Subtypes Versus Non-Cerebellar Healthy Brain Tissue (GTEx)**

*Comparison of gene expression patterns between glioblastoma (GBM), astrocytoma, and oligodendroglioma samples and matched healthy brain data from the GTEx consortium*.

**Supplemental Table 4. Differential Gene Expression Profiles in Pediatric Brain Tumors Versus Non-Cerebellar Healthy Brain Tissue (GTEx)**

*Comparative transcriptomic analysis of pediatric high-grade glioma, diffuse intrinsic pontine glioma (DIPG), low-grade glioma, and ganglioglioma relative to non-cerebellar GTEx brain samples*.

**Supplemental Table 5. Differential Gene Expression Profiles in Medulloblastoma Molecular Subgroups Versus Cerebellar Tissue (GTEx)**

*Analysis of gene expression alterations in medulloblastoma subgroups (Group 3, Group 4, WNT, and SHH) relative to cerebellar samples from the GTEx database*.

**Supplemental Table 6. Differential Gene Expression Profiles in Ependymoma Subtypes Versus Non-Cerebellar Healthy Brain Tissue (GTEx)**

*Transcriptomic comparison of ependymoma subtypes against non-cerebellar healthy brain samples from GTEx*.

**Supplemental Table 7. Differential Gene Expression Profiles in Meningioma Versus Non-Cerebellar Healthy Brain Tissue (GTEx)**

*Comparison of gene expression in meningioma samples with corresponding non-cerebellar healthy brain tissue from GTEx*.

